# Directed neural interactions in whole-brain resting-state fMRI: a comparison between Granger Causality and Effective Connectivity

**DOI:** 10.1101/2024.02.22.581068

**Authors:** Michele Allegra, Matthieu Gilson, Andrea Brovelli

**Author notes:** equal contribution (Matthieu Gilson, Andrea Brovelli).

## Abstract

A key challenge for network neuroscience is to understand the role of interactions between brain regions and how they contribute to the encoding and broadcasting of information within cognitive processes. This requires computational tools to infer directional relations between brain regions from neural time series. For fMRI, the most common approaches are based on Granger causality (GC) analysis and effective connectivity (EC) models. Despite their different conceptual framing, GC and EC models for fMRI are based on similar mathematical assumptions, grounded on continuous- and discrete-time linear stochastic models. Based on a mapping between multivariate Ornstein-Uhlenbeck (MOU) and multivariate autoregressive (MVAR) processes, we analytically obtain an approximately quadratic relation between EC and GC, after rescaling the GC to compensate for unequal noise variances of the source and target. Simulations show that these relations can be observed in finite time series only if a large amount of data is available, implying that they may emerge only at a group level in real fMRI experiments. We verified this prediction by systematically comparing EC and GC in a large-scale fMRI data set from the Human Connectome Project. Overall, our findings can provide methodological and interpretational guidance in the usage of GC and EC for brain network reconstruction by clearly elucidating what is common between the two methods, but also their respective specificities and biases.

## Introduction

A central hypothesis in neuroscience posits that cognitive functions arise from the coordinated activity of neural populations distributed over large-scale brain networks [1, 2] and from the emergence of network-wide and self-organized information routing patterns [3, 4]. Therefore, the estimation of interactions between brain regions is a key challenge for modern neuroscience, and indispensable for understanding the encoding and broadcasting of information relevant for cognitive processes in the brain. IN recent decades, functional magnetic resonance imaging (fMRI) has been increasingly used to study large-scale brain networks, due to its fine spatial resolution [5, 6]. The most widely used approach to identify brain networks from ongoing activity observed by resting-state fMRI (rs-fMRI) [7] is based on functional connectivity (FC), i.e., the correlation of the rs-fMRI signals of pairs of brain areas [8]. Despite its simplicity, FC analysis leads to reproducible and reliable findings [9] about brain network activation during cognitive tasks, alteration due to neuropathologies, and specificity for individuals [10]. It has provided a considerable wealth of information about the large-scale organization of brain activity, its relation to the underlying anatomy, and the neural signatures of individual behavioral traits [10–16].

However, FC methods do not provide information about potential directional or asymmetric interactions between brain areas and are highly sensitive to third-party effects. To bypass these limitations, there exist several methods that exploit the temporal structure of ongoing fMRI signals to capture direct (non-mediated) and directional connections between brain areas [17]. Although a wide array of methods have been proposed in the literature [18], most applications rely on two classes of methods.

The first class is based on on the Wiener-Granger principle [1,19,20], which equates the presence of a directional connection with the capability of a time series to forecast another. The degree of forecasting is quantified information-theoretically, via the transfer entropy [21], or assuming an autoregressive model, via *Granger causality* (GC) [19]. GC is interpreted in terms of influence or information flow between brain regions, and is equivalent to TE for Gaussian variables [22]. Applied to rs-fMRI, most studies assume a first-order multivariate autoregressive (MVAR) model to calculate GC [23–31].

The second class relies on the concept of *effective connectivity* (EC), assuming a generative network model whose dynamics are optimized through the model connectivity to reproduce statistics of empirical data [8]. This concept was widely popularized through dynamic causal modeling (DCM) [32]. To address whole-brain rs-fMRI connectivity, several simplifications of the standard DCM are required, allowing for the analysis of large networks (*n* ≳ 50 areas; *n*^2^ ≳ 2500 connections) in absence of external input (i.e., of events that allow for an easy reconstruction of the hemodynamic response function). The most popular approach, regression DCM (rDCM [33]) assumes linear generative dynamics and a linearized hemodynamic response [34] (for a similar approach, see also [35]). The estimated EC weights are still interpreted as the strength and sign of directional interactions between neuronal populations, as reflected by the corresponding time series. An alternative approach for whole-brain EC modelling is based on the Lyapunov optimization of multivariate Ornstein-Uhlenbeck processes (MOU-EC) [36, 37]. In this approach, hemodynamics is neglected and EC weights are inferred via gradient descent from non-lagged and lagged cross-covariances. In fact, rDCM and MOU-EC mainly differ in terms of inference methodology (Lyapunov optimization vs. variational Bayes), rather than in the underlying generative model. Another popular EC model [38, 39] considers each area as a weakly nonlinear oscillator coupled to other areas through EC, which is estimated by heuristically optimizing model parameters to best reproduce observed cross-correlations. Whole-brain EC is commonly used to uncover sub-networks and hierarchies of brain regions specific to cognitive tasks or pathological conditions, both at the group level or at the single-subject level [40–44].

On one hand, these two classes of methods provide conceptually different metrics. GC is a non-signed measure of *influence*, quantifying the amount of directional interaction between nodes. Instead, EC is a signed metric depicting the effect of a local activity increase on downstream regions, which can be both positive (‘excitatory’) or negative (‘inhibitory’). On the other hand, the two classes of methods are based on similar mathematical assumptions. Although GC is usually seen as data-driven, most applications actually assume a generative model in the form of an MVAR, which is closely related to linear models used for EC estimation.

In this study, we set out to thoroughly discuss the consistency between these two classes of methods in the context of fMRI. Our first main goal was to trace the common mathematical formalism underlying popular approaches for GC and EC in the context of fMRI. This has led us to uncover analytic relations between MVAR-based GC and MOU-based EC. Although these relations are derived for specific versions of GC and EC, and they are expected to hold only in the limit of vanishing sampling noise, they nevertheless provide a clear theoretical framework for discussing GC-EC alignment. The second main goal was to investigate the GC-EC alignment in realistic conditions (in terms of time series length, which determines that amount of sampling noise). This step was accomplished through numerical experiments and served to properly interpret the alginment/misalignment observed in real data. Finally, the third goal was to investigate the EC-GC alignment in real fMRI data. To this end, we leveraged the standard rs-fMRI data provided by the Human Connectome Project [45,46]. In short, our analysis will reveal that a consistency between EC and GC, both in terms of connection strength and connection asymmetry, can be obtained only at a group level, and when properly accounting for unequal node variances and the existence of signed connections. Overall, our analysis is intended to provide methodological guidance in the usage of GC and EC for brain network reconstruction, by clearly elucidating what is common between the two methods, but also their respective specificities and biases.

## Materials and Methods

### Multivariate Ornstein–Uhlenbeck (MOU) and autogressive (MVAR) models

In this work, we set out to discuss the relation between Granger Causality (GC) and Effective Connectivity (EC) approaches to characterize directed interactions in fMRI. Our first step is therefore to show that, under specific assumptions, one can find *analytic relations* between effective connectivity and Granger causality. These relations may hold, up to some approximation, also when the assumptions are relaxed. Therefore, a discussion of these relations should be the starting point if one wishes to correctly frame a systematic comparison between GC and EC.

The core of EC methods is assuming that neural signals are generated by a continuous-time dynamical system with unknown inter-areal couplings. The observed (discrete) time series are assumed to correspond to observations of the continuous system taken with a given sampling time Δ*t* (corresponding to the repetition time for fMRI), and the methods try to reconstruct the couplings from the available data. The resulting estimates are called *effective connections*. Different methods differ depending on the choice of dynamical system and inference method, but a large majority assume a linear dynamical system. Here we focus our attention on MOU-EC method [36], which maximally facilitates discussion of the GC-EC relation. This method assumes that the neural time series **x**(*t*) are produced by a 1^st^ order, linear, Gaussian, stochastic dynamical system, i.e., a multivariate Ornstein-Uhlenbeck process (MOU) given by

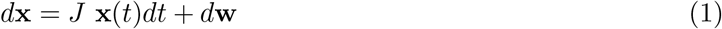

where *J* is the system’s Jacobian and *w* a noise term. The term ‘Jacobian’ is used here because linearization of any dynamical system with noise *d***x** = **f** (**x**(*t*))*dt* + *d***w** will lead to an equation like (1), with *J* = *∂***f** */∂***x** the Jacobian of the vector field **f**. The MOU-EC method assumes that 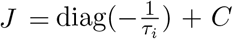, i.e., the Jacobian includes a diagonal term with typically homogeneous or nearly homogeneous time constant *τ*_*i*_ ≈ *τ*, and an off-diagonal connectivity term (*C*). The main role of the diagonal term is to stabilize dynamics, which (in the absence of external input or noise) would produce a relaxation towards a quiescent state (**x** = 0) with a typical time scale *τ*. Even in the presence of noise, *τ* effectively determines the typical autocorrelation time of the dynamics. The off-diagonal terms determine the strength of inter-areal couplings in the model, and they can be both positive (effectively ‘excitatory’ interaction) and negative (effectively ‘inhibitory’ interaction). The noise is assumed to be white, i.e., *d***w** is a Wiener process with covariance matrix ⟨*d***w***d***w**^*T*^ ⟩ = ∑ *dt*.

GC methods assume that the observed (discrete) time series, with sampling time Δ*t*, are generated by a multivariate autoregressive (MVAR) process. In the scope of fMRI, due to the slow sampling frequency, a common choice is using a 1^st^ order multivariate MVAR process

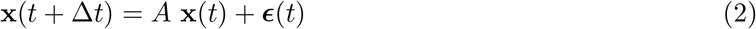

where ***ϵ***(*t*) ~ 𝒩 (0, *S*) is a Gaussian noise with zero mean and covariance matrix *S*. To quantify the Granger-causal effect of node *i* on node *j*, one can measure the relevance of *x*_*i*_(*t*) in predicting *x*_*j*_(*t* + Δ*t*). To this aim, one can define a a “reduced” MVAR process where the influence of node *i* is removed:

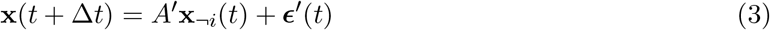

where **x**_¬*i*_(*t*) is obtained from **x**(*t*) by removing its *i*-th component and ***ϵ***′(*t*) is Gaussian with zero mean and covariance matrix *S*′. The Granger causal effect of node *i* on node *j* is given by the log-ratio of the variances,

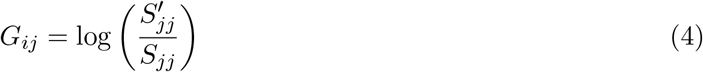

Note that we consider the conditional version of GC, which means that the linear regression in Eq. (3) includes all remaining nodes in the network. To compute GC, one should estimate the variances 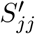 and 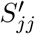 from the observed time series. In addition ((4)), Geweke [47] defined the ‘instantaneous’ Granger causality *I*_*ij*_, a non-directional connectivity metric that measures the correlations in the innovations *ϵ*_*i*_ and *ϵ*_*j*_:

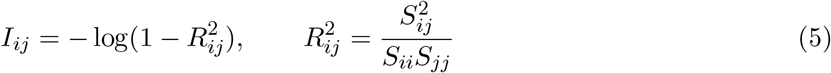

The name ‘instantaneous’ comes from the fact that, from the point of view of the observer, these correlations cannot be predicted autoregressively, i.e., from past values of the time-series, and thus appear as instantaneous. Although the instantaneous causality is often discarded in Granger causality analysis, it may capture a large part of the interdependence between two time series.

As linear-feedback dynamic systems, the Ornstein-Uhlenbeck process and the 1^st^ order MVAR rely on mutually consistent assumptions about the nature of the underlying network process, only differing by the continuous vs. discrete nature of time. As shown in the Appendix, the respective connectivity terms (*J* and *A*) can be formally related, and this translates into a mathematical relationship between Granger causality estimated from the MVAR and the EC estimated from the MOU.

### Regression DCM

For resting-state fMRI, alongside MOU-EC, one of the most used methods to infer EC is Regression DCM [33, 48]. This is a computationally efficient variant of DCM designed for whole-brain network analysis. Briefly, rDCM incorporates several key modifications to the standard DCM framework that enable large-scale connectivity estimation involving the whole brain. The approach distinguishes between the neural time series **x**(*t*) and the observed BOLD signal **y**(*t*), which is assumed to be related to the neural signal through convolution with a hemodynamic filter. rDCM assumes a linear dynamic model and a linearized observation model,

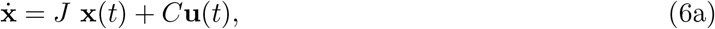

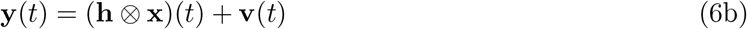

where ⊗ denotes a convolution, **u**(*t*) is external input, **h** is a (linearized) hemodynamic filter, and **v** is Gaussian observation noise. Note that **y** is observed only at discrete times *t, t* + Δ*t, t* + 2Δ*t*, etc. When modeling ongoing activity, the external input is absent, **u**(*t*) = 0. Passing to the frequency domain and discretizing, the system becomes

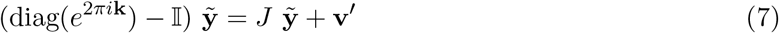

where 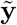 denotes the discrete Fourier transform, 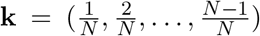, and **v**′ is an (in principle correlated) Gaussian noise term. The simplifying assumption is made that noise affecting different regions is independent, **v**′ ~ 𝒩 (0, diag(***ρ***^2^)) where ***ρ*** is the vector of noise variances.

Given Eq. (7), the estimation of *J* can proceed via Bayesian linear regression. The assumption of independent noise inputs translates into the fact that the posterior factorizes into the product of independent terms for afferent connections to each region, allowing for efficient inference. Note that Eq. (7) is completely independent of **h** (this would not hold in the presence of external inputs, **u**(*t*)≠ 0). In other words, we could safely replace **h** with the identity filter, which would reduce the system to

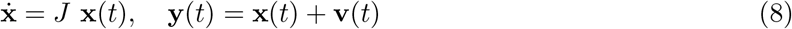

which brings it in a form very similar to (1). Regression DCM is implemented in the TAPAS toolbox for Matlab [49], but a Python implementation is also available [50].

### Network simulations

The relations (9), which imply a piecewise monotonic relation, and hence a perfect alignment, between GC and IC, are theoretical: they hold between the model connectivity *C* and *G, I* computed on the true (population) covariance matrix. In finite data, these relations should reflect in analogous relations between the effective connectivity *Ĉ* reconstructed from the data and the estimated Granger causality measures 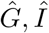 computed from empirical (sample) covariance matrices. Goal #2 of our study is assessing whether, and to what extent, the predicted relations hold in finite-length time series. We generated time series using the MOU model (1) with process time scales *τ* ∈ [0.1, 10] and a fixed sampling period Δ*t* = 1. The model connectivity *C* was a random matrix with probability *p*_1_ = 30% of connection between each pair of nodes. We considered networks of *N* = 10, *N* = 40 and *N* = 100 nodes, The weights were drawn from a Pareto distribution 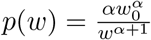 with *α* = 5 and *w*_0_ = 0.1*/τ* (for *N* = 5); with *α* = 3 and *w*_0_ = 0.1*/τ* (for *N* = 40); with *α* = 3 and *w*_0_ = 0.5*/τ* (for *N* = 100). With a probability *p*_2_ = 30%, a pair of reciprocal connections (*C*_*ij*_,*C*_*ji*_) was chosen and their sign was flipped. Thus, roughly 30% of connections are negative. Finally, for all pairs of non-zero reciprocal links, a random number 0 *< r <* 0.2 was extracted, and one of the two connections was multiplied by 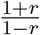, generating asymmetries in reciprocal connections.

In all simulations, we used a diagonal noise matrix 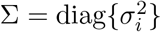. We made the values of *σ*_*i*_ dependent on *τ*, aligning the noise time scale and the process time scale (stated otherwise, to align the magnitude of ∑ and that of *C*). We generated random numbers 0.2 ≤ *s*_*i*_ ≤ 5, and considered 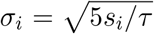, i.e., different nodes were affected by noise of different magnitude, with noise magnitudes spanning roughly an order of magnitude.

### Human resting-state fMRI data

Goal #3 of our study was to investigate relations between effective connectivity and Granger causality in real fMRI data and test predictions previously obtained through analytical derivations and simulation results. We used the subset of 100 unrelated subjects (54 females, 46 males) from the Human Connectome Project (HCP) [45]. For the main analysis, we used the left-right (LR) phase-encoding runs from the first session resting state fMRI data. We later replicated the analysis for the right-left (RL) phase-encoding runs. Time series had 1200 time points with a TR of 0.72 sec, comprising ≈ 15 minutes of scanning. The full description of the imaging parameters and minimal preprocessing pipeline is given by Ref. [51]. In short, after correction for motion, gradient, and susceptibility distortions the fMRI data was aligned to an anatomical image. The aligned functional image was corrected for intensity bias, demeaned, and projected to a common surface space, which resulted in a cifti-file. Artifacts were removed through Independent Component Analysis (ICA) using the FSL’s MELODIC tool paired with the FMRIB’S ICA-based X-noisefilter. No additional global signal regression was applied. All fMRI data were filtered between 0.01 and 0.1 Hz to retain the relevant frequency range for further analysis of the BOLD signal. Functional data can be mapped to different spatial resolutions using the Schaefer parcellation [52], which optimizes local gradient and global similarity measures of the fMRI signals. Here, we selected the parcellation consisting of 100 regions, each being linked to one of the 7 networks: visual (VN), somatosensory (SMN), dorsal attention (DAN), ventral attention (DAN), limbic (LIM), xxx (CN), default-mode network (DMN). For both fMRI datasets, regional time series were extracted using Workbench Command provided by the HCP.

### Statistical analysis of directional connections

For each participant, we computed 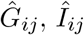 using the covariance-based approach [28, 53] and *Ĉ*_*ij*_ using the Lyapunov optimization method [36]. For each individual connection (in 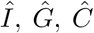, *Ĉ*), we estimated a group-level significance using a t test. Connections were considered significantly positive if *T >* 0 and *p <* 0.05, significantly negative if *T <* 0 and *p <* 0.05, non-significant if *p >* 0.05. P-values were corrected for 9900 multiple comparisons using the false discovery rate approach [54]. For each pair of reciprocal connections (*i* → *j, j* → *i*) we computed connection asymmetries as Δ*Ĉ* = *C*_*ij*_ − *C*_*ji*_, Δ|*Ĉ*| = |*C*|_*ij*_ − |*C*|_*ji*_, 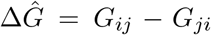. We estimated the group-level significance of connection asymmetry using a t test. Asymmetry was considered significant in the direction *i* → *j* if *T >* 0 and *p <* 0.05, significant in the direction *j* → *i* if *T <* 0 and *p <* 0.05, non significant if *p <* 0.05. P-values were corrected for 4950 multiple comparisons using the false discovery rate approach [54].

## Results

### Theoretical relations between MOU connectivity and Granger causality

As linear-feedback dynamic systems, the Ornstein-Uhlenbeck process and the 1^st^ order MVAR rely on mutually consistent assumptions about the nature of the underlying network process, only differing by the continuous vs. discrete nature of time. As shown in S.I. Appendix, the respective connectivity terms (*J* and *A*) can be formally related, and this translates into a mathematical relationship between Granger causality estimated from the MVAR and the EC estimated from the MOU. Given that the MOU and the MVAR are related by a (non-infinitesimal) discretization step, their relation is critically modulated by the relation between the timescale of the MOU process (*τ*) and that of the sampling (Δ*t*). We can distinguish three regimes: *τ* ≪ Δ*t, fast* dynamics; *τ* ≈ Δ*t, matched* dynamics; *τ* ≫ Δ*t, slow* dynamics. To understand the difference between these regimes, one should consider that in a MOU process the influence of *x*_*j*_(*t*) is felt by *x*_*i*_(*t*′), with *t*′ *> t*, only for *t*′ ≲ *τ*. Therefore, one can directly observe this influence only if Δ*t < τ* (slow dynamics). If Δ*t > τ* (fast dynamics), one can only observe a sort of ‘time-integrated’ effect of the reciprocal influences between *x*_*i*_ and *x*_*j*_, which appears as a non-directional feedback between the two time series. In the Granger-Geweke formalisms, this is captured the instantaneous causality. Our theoretical analysis (S1 Text) leads to a relationship that is valid in the limit of weak coupling for slow dynamics and (to a certain extent) for matched dynamics, but not necessarily for fast dynamics. For those regimes, the analytical relation is given by:

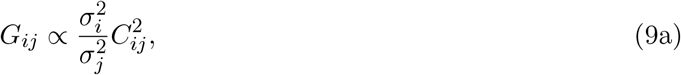

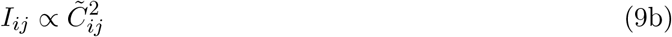

Here, *σ*_*i*_ = ∑_*ii*_ is the noise variance of node *i*; 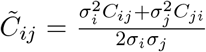 is a mean of the reciprocal Effective Connectivity weights between *i* and *j*, weighted by the noise variances *C*_*ij*_; *G*_*ij*_ is Granger causality; and *I*_*ij*_ is the instantaneous (Granger) causality” (IC)

If the input noise is homogeneous across all nodes (*σ*_*i*_ = *σ*), these relations simplify as:

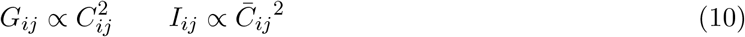

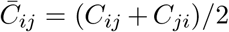 being the symmetrized effective connectivity. These relations mean that standard and instantaneous Granger causality provide an estimate of the *square* of the node coupling in a MOU process. In the general case of inhomogeneous noise variance, we can still retrieve an approximately quadratic relation. Since, approximately, 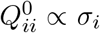, where *Q*^0^ = ⟨**xx**^*T*^ ⟩ is the covariance matrix of **x**, we can introduce “corrected” versions of *G* and *I*,

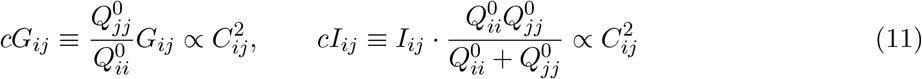

Finally, the ratio between Granger causality and instantaneous causality is approximated as:

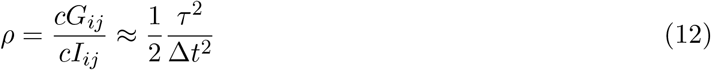

This means that for slow dynamics *cG* ≫ *cI*, whereas *cG* ≪ *cI* for fast dynamics.

Thus, assuming that a continuous MOU model is a valid generative model for the observed time series, the information about the model connectivity *C* is differentially captured by *G* and *I* depending on the sampling regime. When the dynamics is faster than sampling, the values of the observed time series at the previous time point are poorly predictive of the next time point. Thus, regardless of the strength of effective connections, GC remains low, in line with previous work [55]. The presence of strong effective connections still creates a strong statistical dependency between the two time series, but most of this interdependence appears as ‘instantaneous’ correlation that cannot be predicted on the basis of previous time points.

### Validation of the theoretical relation in simulated data

To test the analytical relations derived in the previous sections, we generated a MOU process with known (“ground truth”) connectivity and tested the above quadratic relations between the model connectivity (*C* in equations) and Granger causality measures (*G* and *I*, respectively). We considered a network of *N* = 40 nodes, with probability *p* = 0.3 or a directional link between each pair of nodes. We randomly selected link weights from a power-law distribution to reproduce the heavy-tailed weight distribution observed in typical brain networks, and we allowed for connectivity asymmetries and negative weights (see Methods for details). We also considered the case of variable noise across nodes (spanning more than one order of magnitude). We fixed the sampling time Δ*t* = 1 and considered different values of *τ* to explore all the three dynamical regimes (fast, matched and slow). The results are shown in Fig. 1b. We compared the consistency of *C* with the uncorrected (*G, I*) and corrected (*cG* and *cI*) versions of Granger causality. The quadratic relations were satisfied to a very good accuracy for the corrected case. When the correction was not applied, the quadratic relation between *C* and *G* was degraded. Furthermore, as expected, we observed that *I* prevailed over *G* in the fast regime, while the opposite occurs in the slow one. Although the theoretical relations were derived for the conditional versions of *G* and *I*, very similar results were obtained by considering the respective unconditional versions (Fig. S1).

**Figure 1:**
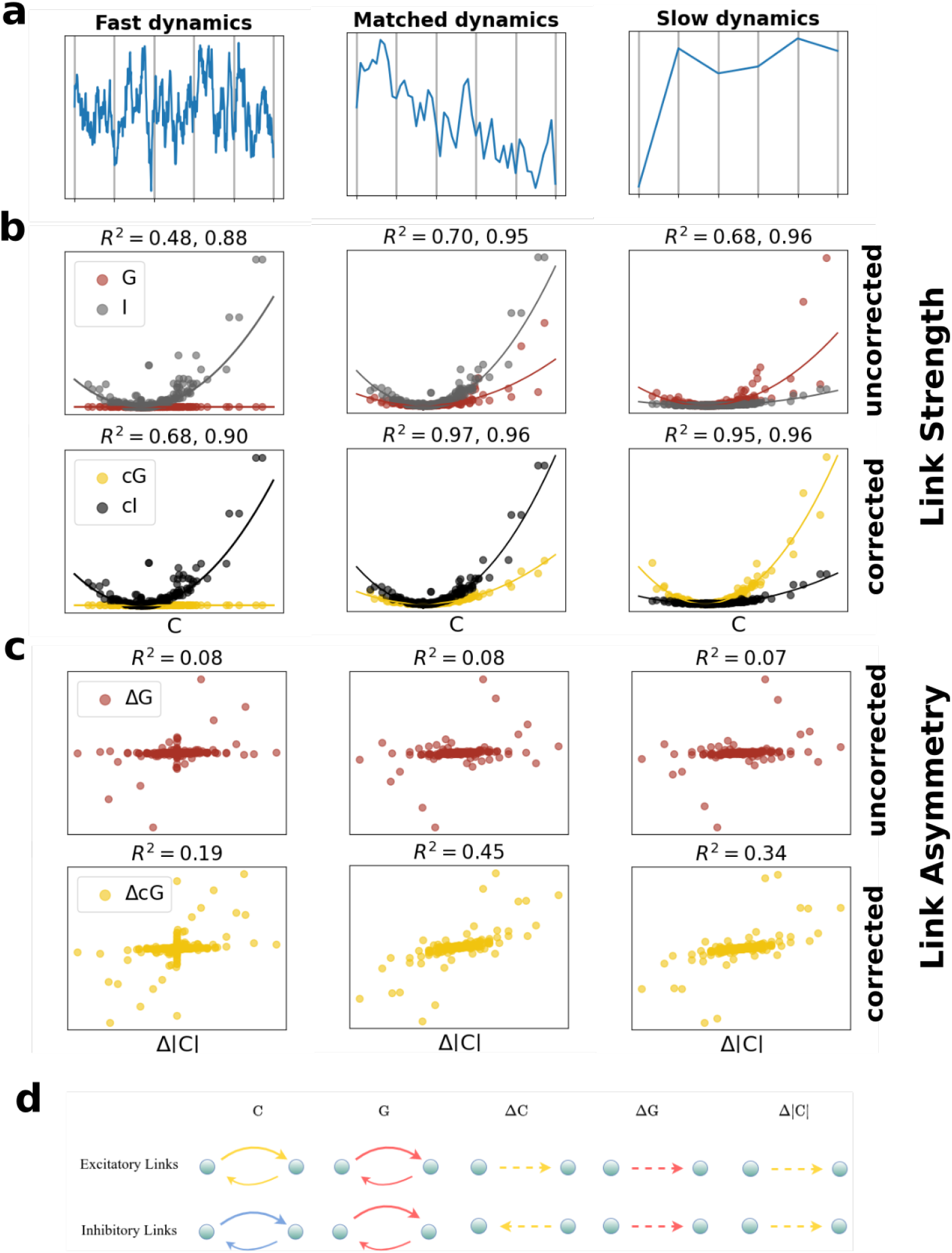
Theoretical relations between model connectivity and Granger causality measures. We considered a random network of *N* = 40 nodes evolving according to the MOU dynamics (1). The corresponding MVAR process has a sampling time Δ*t* = 1. **(a)** Depending on the characteristic time *τ* of the MOU, we obtained three regimes: slow (*τ* = 0.1), matched (*τ* = 1) and fast (*τ* = 10). **(b**) Relation between the model weights *C* and the corresponding values of *G* (in red) and *I* (in black) for each connection, without correcting for unequal input noise variances (top row) and correcting (bottom row). Each dot corresponds to a pair (*i, j*), and solid lines to the predicted scalings. **(c**) Relation between the asymmetry in (the absolute strength of) the model weights Δ|*C*| and the corresponding asymmetries in Δ*G* (in red), without correcting for unequal input noise variances (top row) and correcting (middle row). Each dot corresponds to a pair (*i, j*). **(d**) Conceptual relation between Δ*G*, Δ*C* and Δ|*C*| for the three cases in which *C*_*ij*_, *C*_*ji*_ are positive (‘excitation’) or negative (‘inhibition’).

The theoretical relation suggests that *G* could also provide information about the link asymmetries in the strength of reciprocal connections *C*_*ij*_ and *C*_*ji*_. In other words, when considering asymmetries

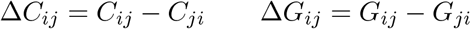

one would expect a relation between Δ*G*_*ij*_ and Δ*C*_*ij*_. This intuition is correct, but there are important caveats. Firstly, the quadratic relation (11) implies an relation for asymmetries only if the noise variances coincide, *σ*_*i*_ = *σ*_*j*_. Therefore, correcting for unequal noise variances becomes truly essential to get concordance. Secondly, the relation (11) is quadratic (not monotonic), and implies that *G* is insensitive to whether the underlying connections are ‘excitatory’ (*C*_*ij*_ *>* 0) or ‘inhibitory’ (*C*_*ij*_ *<* 0). If all connections were excitatory, Δ*G*_*ij*_ and Δ*C*_*ij*_ would always be concordant. However, when effective connections are signed, this is not necessarily the case. This point is exemplified in Fig. 1d. A positive value of Δ*C*_*ij*_ *>* 0 associated to a net asymmetry *i* → *j* of the effective connection can correspond to different situations, depending on the sign of *C*_*ij*_, *C*_*ji*_:

- *C*_*ij*_ *>* 0, *C*_*ji*_ *>* 0, Δ*C*_*ij*_ *>* 0: both connections are excitatory, and the influence of *i* on *j* is larger than the reverse. Δ*C*_*ij*_ and Δ*G*_*ij*_ are concordant.
- *C*_*ij*_ *<* 0, *C*_*ji*_ *<* 0, Δ*C*_*ij*_ *>* 0: both connections are inhibitory, but in this case it is the influence of *j* on *i* that prevails. Δ*C*_*ij*_ and Δ*G*_*ij*_ are discordant.
- *C*_*ij*_ *>* 0, *C*_*ji*_ *<* 0, Δ*C*_*ij*_ *>* 0: one connection is excitatory and the other inhibitory. In this case, Δ*C*_*ij*_ and Δ*G*_*ij*_ are not necessarily concordant; Δ*G*_*ij*_ could even be zero. This case is quite rare in real data (see the following section).

To retrieve an alignment between *C* and *G* one should compare the net influence asymmetry measured by Δ*G* with a net difference in the *absolute strength* (positive or negative) of effective connections, measured by

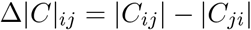

In Fig. 1c) we show the consistency of Δ|*C*| and Δ*G*, Δ*cG*. We observe an alignment between Δ|*C*| and Δ*cG* only in the matched and fast regimes, where interaction unfolds over a time scale longer than the sampling time, and interaction asymmetries are more easily measurable.

### Robustness of the theoretical relation with respect to sampling noise

In the previous section, we have shown that when data are generated by a continuous, linear system with Gaussian noise (i.e., a MOU process), Granger causality is related to the model connectivity by a quadratic relation. In real data, however, this relation may be hidden under sampling noise, as (i) Granger causality must be estimated from finite samples; we will denote by 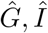 the resulting estimates, which will differ from the theoretical values *G, I* obtained in the infinite sample limit (ii) the model connectivity *C* is generally unknown; effective connectivity *Ĉ* provides an estimate of it. To test the effects of sampling noise, we considered simulations of the same model as in the previous section. We varied the characteristic time of the MOU model, which determines the dynamical regime, and the simulation length ℒ, which determines the amount of sampling noise. We estimated GC using the covariance-based approach, which does not try to fit an MVAR but computes GC directly from the non-lagged and lagged covariance matrices 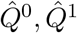. EC is estimated using two approaches, rDCM and MOU-EC. In this way, we can guarantee that conclusions are not contingent on the usage of a specific method (MOU-EC).

We first tested if relations between the model connectivity and Granger causality would also hold when considering sampling noise affecting Granger causality.

To this end we performed two fits: (i) a quadratic fit of the model connectivity *C* on 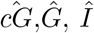, to test the predicted quadratic relations for link strengths (ii) a linear fit of Δ|*C*| on 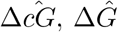, to test the predicted linear relations for link asymmetries. The goodness of fit was quantified through the *R* coefficient (*R*^2^ → 1 corresponding to a perfect fit). The results are shown in Fig. 2a,b.

**Figure 2:**
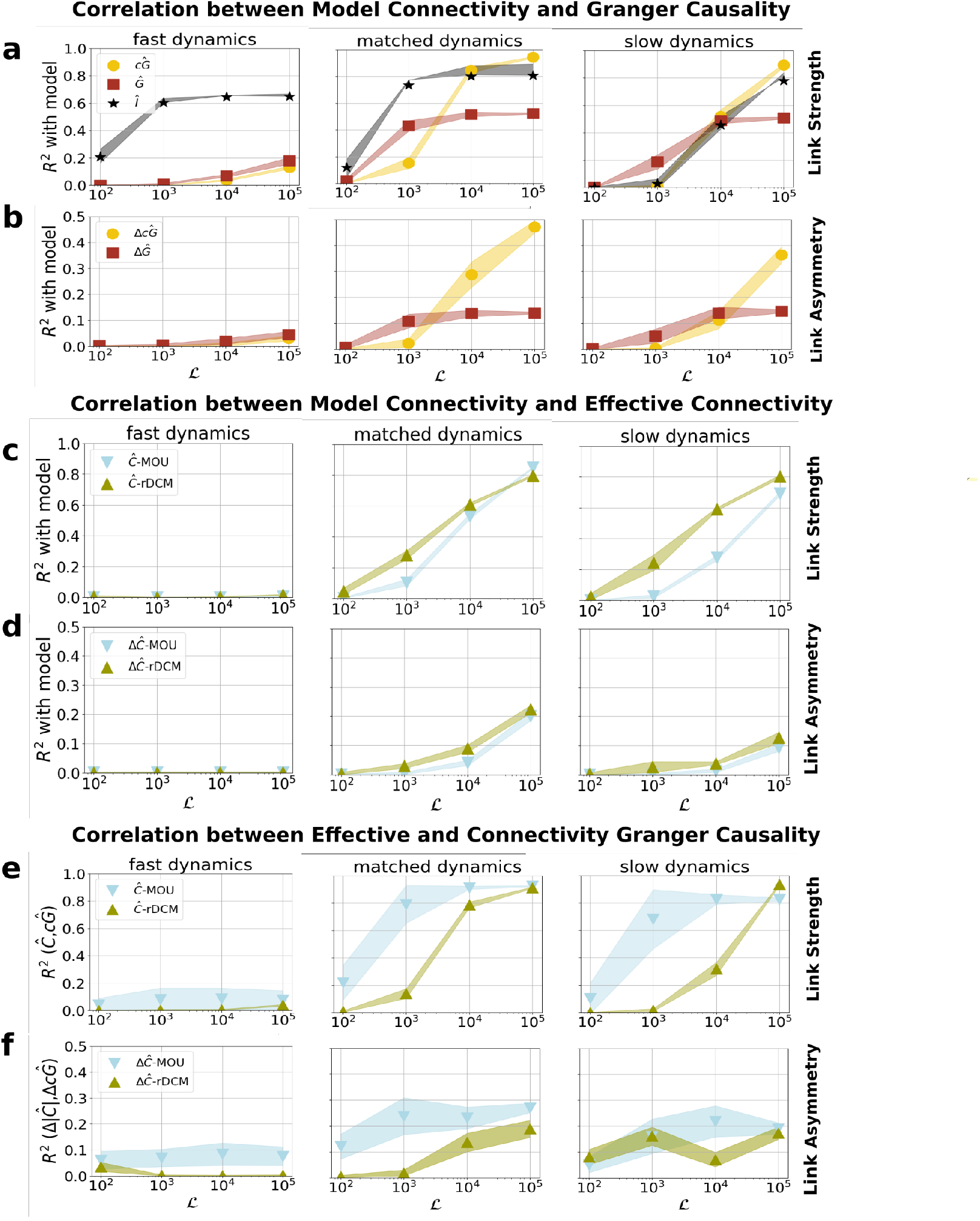
Relations between model connectivity, effective connectivity, and Granger causality measures for finite sampling time. A random network was simulated for different durations ℒ and different dynamical regimes: fast (*τ* = 0.1), matched (*τ* = 1) and slow (*τ* = 10). **(a)** Variance of the model connectivity *C* explained by uncorrected 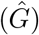 and corrected 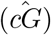 Granger causality, and instantaneous Granger causality 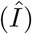. **(b)** Same as (a), but considering asymmetry in the absolute link strength Δ |*C*| vs 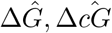. **(c)** Variance of the model connectivity *C* explained effective connectivity estimated from the dat via MOU-EC (*Ĉ*-MOU) and rDCM (*Ĉ*-rDCM), **(d)** Same as (c), but considering asymmetry in the absolute link strength Δ|*C*| vs Δ*Ĉ*-MOU,Δ*Ĉ*-rDCM. **(e)** Variance of the effective connectivity (estimated with MOU and rDCM) explained by corrected Granger causality **(f)** Same as (e), but considering link asymmetries Δ|*Ĉ*|-MOU, Δ*Ĉ*-rDCM vs 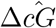.

For *fast* dynamics, 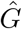 and 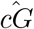 are poorly related to *C*, consistent with the fact that one cannot appreciate the directionality of interactions in this regime. Correspondingly, there is also no correlation between link asymmetries in 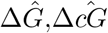 and Δ|*C*|. Instead, the instantaneous causality *I* reaches high values of *R*^2^ ≈ 0.6 as long as the simulation is sufficiently long (ℒ ≳ 10^3^). For *matched* dynamics, we observed a strong alignment between 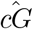 and *C*^2^ (*R*^2^ *>* 0.8) but only for very long sampling length (ℒ ≳ 10^4^). Alignment between 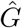 and *C*^2^ was better (*R*^2^ ≈ 0.4) for ℒ = 10^3^ (a quite realistic length for fMRI), but did not improve for long sampling lengths. In terms of asymmetry we obtained a similar picture, as 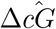 aligned with Δ|*C*| but only for very long sampling lengths, reaching *R*^2^ ≈ 0.5 for ℒ = 10^5^. Instantaneous causality aligned well with *C*^2^ for ℒ ≳ 10^3^. Finally, for *slow* dynamics, results were similar to those obtained for matched dynamics, but the values of *R* decreased, especially for the instantaneous causality that was completely uncorrelated to *C* for a realistic sampling length (ℒ = 10^3^. Overall, the predicted relations between model connectivity and Granger causality, in terms of link strengths and asymmetries, were best observable when the characteristic time of the MOU process corresponded to the sampling time (matched dynamics); the relation between *C* and *I* was robust to sampling noise, while the relation between *C* and *cG* (and between Δ*C* and Δ*cG*) was not, emerging only for long sampling lengths.

Secondly, we asked whether relations would still hold when replacing the model connectivity *C* with effective connectivity estimates *Ĉ*. To this aim, we first performed a general assessment of the quality of estimates obtained with MOU-EC and rDCM, by performing a linear fit of *C* on *Ĉ*. Results are shown in Fig. 2c-d. For fast dynamics, estimates are completely unreliable, remaining around *R*^2^ ≈ 0 for all sampling lengths, consistent with the fact that directional interactions cannot be properly observed in this regime. For matched and slow dynamics, good estimates generally required large sampling length (ℒ ≳ 10^4^). For link strenghts, the maximum alignment was *R*^2^ ≈ 0.8, for link asymmetry *R*^2^ ≈ 0.2. RDCM yielded more accurate estimates than MOU for realistic sampling lengths (ℒ = 10^3^) and slow dynamics. From these results, we might expect a clear alignment between effective connectivity and Granger causality only if (i) the sampling lengths is large, or (ii) estimates are affected by common biases (e.g., note that covariance based GC and MOU-EC both employ the zero-lag and lagged empirical covariance matrices). We tested this by performing (i) a quadratic fit of effective connectivity *Ĉ* on 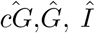 (ii) a linear fit of Δ|*Ĉ*| on 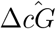. The results are shown in Fig. 2e,f. For fast dynamics, Granger causality and DCM-*C* are very weakly correlated. Unsurprisingly, for fast dynamics we obtained very weak correlations for both link strength and asymmetry. For matched and slow dynamics, we generally obtained a strong alignment between 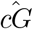 and *Ĉ*-MOU for all ℒ ≳ 10^3^, reaching *R*^2^ ≈ 0.9, even larger than the correlation between model connectivity and effective connectivity. The same occurred for link asymmetry, which reached *R*^2^ ≈ 0.3 (compared to *R*^2^ ≈ 0.2 for model and effective connectivity). Instead, *Ĉ*-rDCM aligned well with 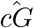 only for a large sampling length (ℒ ≳ 10^4^).

### EC and GC alignment in rs-fMRI data

We considered human resting-state fMRI data of 100 unrelated participants from the Human Connectome Project [45]. For each participant, two recording sessions of 1200 time points (TR=0.7) were available. We mainly considered the first session, using the second session for validation. Upon surface projection and minimal preprocessing, time series were projected onto the Schaefer-100 atlas (100 regions). Regions were divided into seven resting state networks, (RSNs) according to the well-established division proposed by Yeo et al. [56]. Effective connectivity *Ĉ*, that was estimated individually for each participant using MOU-EC and rDCM. Granger causality was estimated individually using the covariance-based approach.

#### Effective connectivity analysis

At a group level, MOU-*Ĉ* featured 3804 significant connections (two-tailed T test over subjects, *p <* 0.01, FDR corrected for 9900 multiple comparisons), among which we found both significantly positive (64%) and significantly negative connections (36%) (Fig. 3a). Reciprocal connections (*i* → *j* and *j* → *i*), tended to be either unidirectional (i.e., when one connection was significant the reciprocal one was non-significant), concordant (both connections were significant and had the same sign; Fig. 3c), or both non-significant. We found only ≈ 1% of pairs of reciprocal connections with discordant sign. Positive-positive reciprocal connections were frequently found within different areas of the same resting state networks (RSNs), both within and across hemispheres. Connections between nodes of different RSNs were predominantly unidirectional (64%). Negative connections were nearly exclusively found between different RSNs (S3 a).

**Figure 3:**
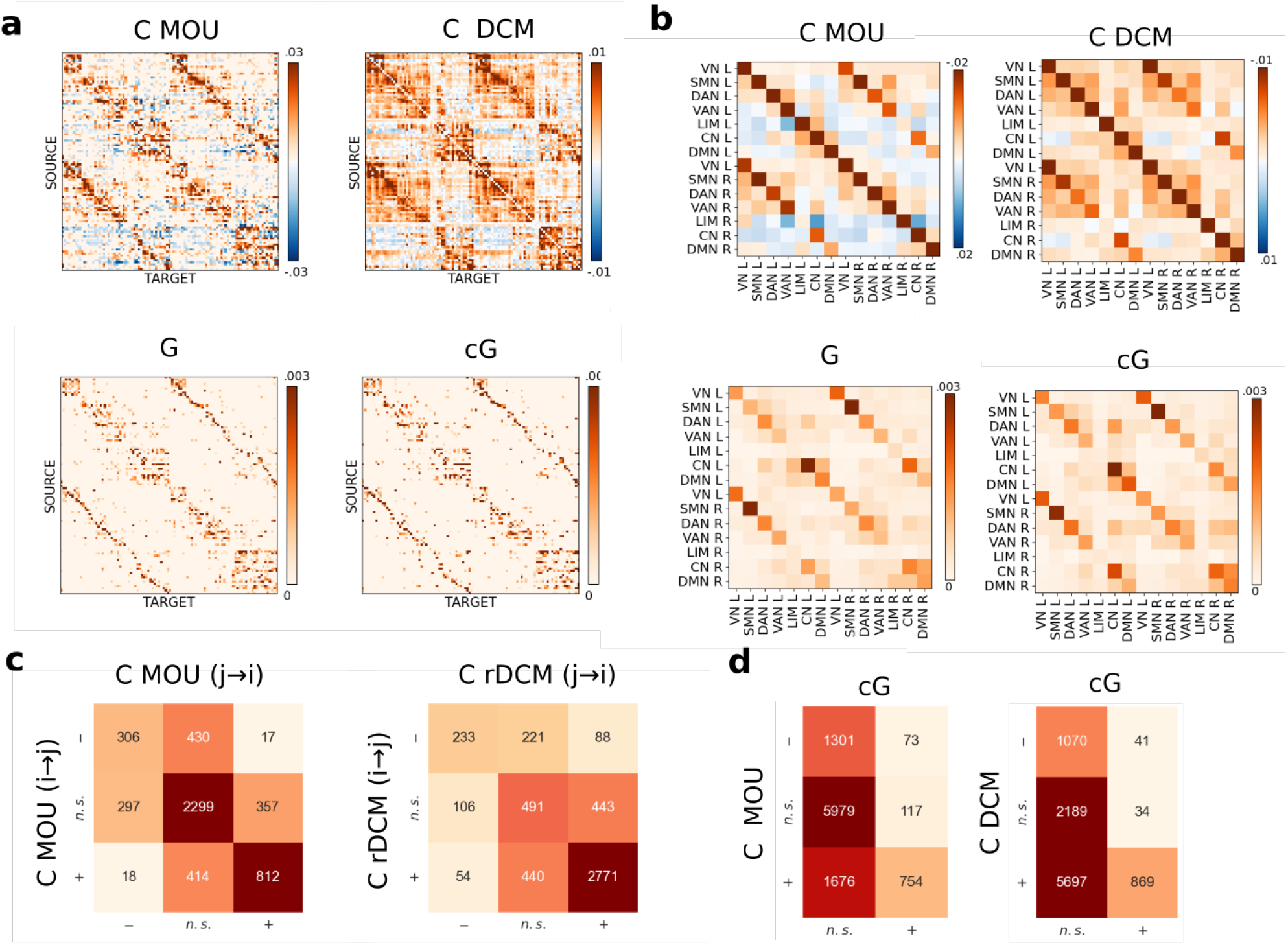
Group-level results of Granger Causality and Effective connectivity in HCP. **(a)** Group-level (average) matrices of 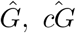, *Ĉ*-MOU, *Ĉ*-rDCM. Only links that are significantly different from zero are shown (T test across subjects, *p <* 0.05 FDR-corrected), in red for positive weights and blue for negative weights. **(b)** Network-wise group-level matrices averaged in 7 functional networks: visual (VN), somatosensory (SMN), dorsal attention (DAN), ventral attention (DAN), limbic (LIM), control (CN), default-mode network (DMN). **(c)** For all pairs of reciprocal connections (*i*→ *j* and *j* →*i*), we counted the number of pairs for which connections are significantly positive (+), significantly negative (−) or non-significant (n.s.) for *Ĉ*-MOU and *Ĉ*-DCM **(d)** We counted the number of connections that are significantly positive/negative or non-significant (n.s.) for *Ĉ* (MOU and rDCM) and and 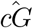.

For all pairs of reciprocal connections, we tested for significant connection asymmetry, Δ*C*_*ij*_ = *C*_*ij*_ −*C*_*ji*_ (T test over subjects, FDR corrected for 4950 multiple comparisons). We found a significant asymmetry for 27% of link pairs (S3 b). Asymmetries between positive connections were found within areas of the same resting state network (RSNs), while asymmetries between inter-RSN links were mostly due to one connection being positive/negative and the reciprocal one being non-significant (S3 b). Only a negligible fraction (2%) of significant asymmetries involved discordant connections.

The estimated rDCM-*Ĉ* showed a much larger number (7676) of significant connections at the group level, and a much lower proportion of negative links (83% positive, 17% negative connections). Reciprocal connections tended to be either unidirectional or concordant; most of them were concordantly positive (Fig. 3c). Roughly 3% of the pairs of significant connections had a discordant sign. Within-RSN connections were almost exclusively positive. Negative connections were almost exclusively found between different RSNs (S3 c). We found a significant asymmetry for the majority (58%) of link pairs (S3C). Asymmetries between positive connections were found both within the same and across different RSNs. Asymmetries between negative connections were only found across different RSNs (S3 d). We averaged *Ĉ* values over the nodes belonging to each of seven resting state networks (RSNs) for both hemispheres, therefore obtaining 14 × 14 network-wise effective connectivity matrices. In Fig. 3c we show significant positive and negative network-wise links (T test over subjects, *p <* 0.01 FDR corrected for 182 multiple comparisons). We observed strong positive connections within each RSN, both within the same hemisphere and across the two hemispheres. On the other hand, negative connections were found between different RSNs, and were strongest for outgoing connections from the control network (CON), limbic network (LIM), and default mode network (DMN). Positive connections were mostly observed within RSNs. For rDCM-*Ĉ*, we mostly observed positive connections. The strongest connections were observed within each RSN. We also see that the RSNs split in two blocks, one formed by ‘task-positive’ networks (visual, somato-motor, dorsal and ventral attention) and the other by ‘task-negative’ network (control, limbic and default mode network). Within-block connections were positive, between-block ones null or slightly negative.

#### Granger causality analysis

Next, we estimated Granger causality. Since the noise variances affecting different nodes were found to be widely different among nodes, spanning roughly an order of magnitude (S4), we expect potential differences between the corrected and non-corrected versions of 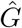. At the group level, the estimated 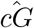 had only 944 significant connections (Fig. 3B) (T test over subjects, *p <* 0.01 FDR corrected for 9900 multiple comparisons). Of these, the overwhelming majority were identified as significantly positive connections by both *Ĉ*-MOU and *Ĉ*-DCM (respectively, 754 and 869). Less than 10% were associated with negative connections in *Ĉ*-MOU and *Ĉ*-DCM (respectively, 73 and 41) (Fig. 3d). The strongest connections were found between homologous regions of the left and right hemishperes. We averaged the 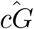 values over the LH/RH nodes belonging to each of seven resting state networks (RSNs), obtaining 14 × 14 network-wise Granger causality matrices. In Fig. 3d) we show significant network-wise links (T test over subjects, *p <* 0.01 FDR corrected for 182 multiple comparisons). The strongest links were again observed within the LH and RH parts of each RSN. We tested pairs of reciprocal links for connection asymmetry: only a small number of link pairs (222, ~5%) presented a significant asymmetry. Of these, most also presented a significant asymmetry in terms of *Ĉ*-MOU and (respectively, 171 and 160). Overall, 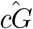 appears much more conservative than *Ĉ* in detecting significant connections and significant asymmetries at a group level. Results for the non-corrected 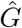 are nearly equivalent.

#### Comparison between effective connectivity and Granger causality

Having analyzed EC and GC separately, we now proceed to a systematic comparison between the two methods. MOU-EC estimates the process time as *τ* = 2.3 ± 0.6 (mean ± st.dev. over subjects). This suggests that fMRI data are approximately in the ‘matched dynamics’ regime. Based on the simulation results, we thus expect to observe an alignment between *C*-MOU and *cG*, and, to a weaker extent, between *C*-rDCM and *cG*. The alignment should become tighter as the sampling noise is reduced. The length of fMRI recordings is fixed (~ 10^3^ time points per participant), but an effective way to reduce sampling noise is to combine data of different subjects to produce group averages. Thus, we expect to see a tighter alignment when averaging *Ĉ*, 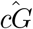 estimates over an increasing number of participants. We considered subgroups of participants of increasing size *n*. For each *n*, we considered *M* = 100 randomly selected subgroups of size *n*. For each subgroup, we averaged *Ĉ* and 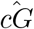 and computed the *R*^2^ between 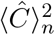 and 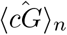 (here, ⟨·⟩_*n*_ indicates a subgroup average over *n* participants). We finally averaged the resulting *R* over the *M* random subgroup choices.

The results are shown in Fig. 4a for *Ĉ*-MOU: *R*^2^ increased rapidly with *n*, reaching values *R*^2^ ≳ 0.6 for *n >* 20. Considering the whole group (*n* = 100), we obtained *R*^2^ = 0.72. Fig. 4a also shows the scatter plot of *Ĉ* and 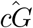, averaged over the whole group of *n* = 100 subjects.

**Figure 4:**
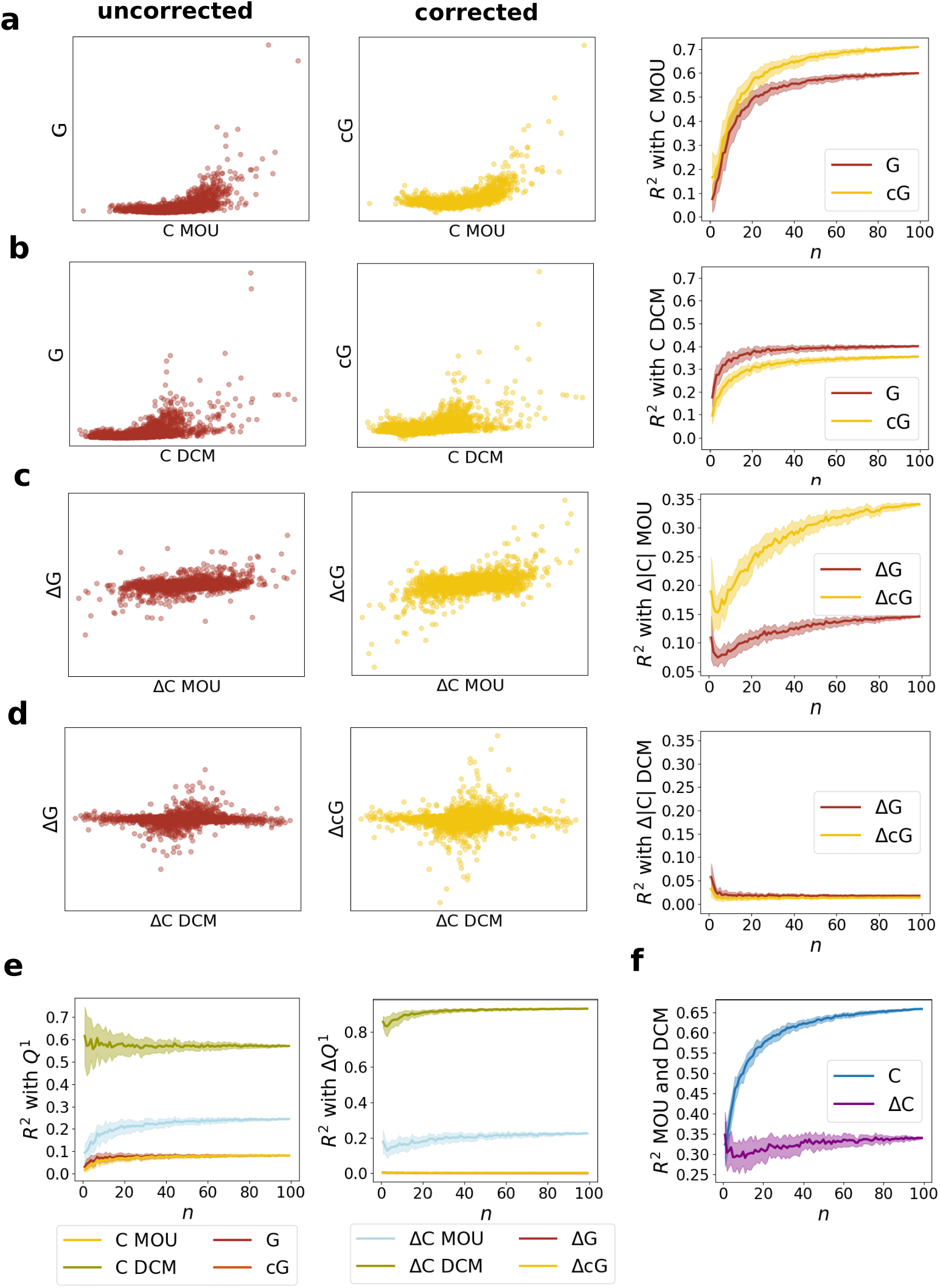
**(a)** effective connectivity *Ĉ* (group average over 100 subjects) vs corrected Granger causality 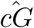 (group average). **(b)** squared Pearson correlation *R*^2^ between group *Ĉ* and group 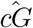 for groups of increasing size *n*. **(c)** effective connectivity *Ĉ* (group average over 100 subjects) vs corrected instantaneous Granger causality 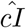 (group average). **(d)** squared Pearson correlation *R*^2^ between group *Ĉ* and group 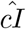 for groups of increasing size *n*. **(e)** asymmetry in effective connectivity Δ*Ĉ* (group average over 100 subjects) vs asymmetry in corrected Granger causality 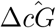 (group average). **(f)** squared Pearson correlation *R*^2^ between group Δ*Ĉ* and group 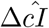 for groups of increasing size *n*.

When considering the non-corrected version of Granger causality 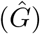, we obtained values of values *R*^2^ ≳ 0.5 for *n >* 20, up to *R*^2^ = 0.6 for *n* = 100. In Fig. 4b, we show results for *Ĉ*-rDCM: *R*^2^ again increases with *n*, reaching values *R*^2^ ≳ 0.3 for *n >* 20. Considering the whole group (*n* = 100), we obtained *R*^2^ = 0.35. The alignment with the non-corrected 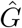 was slightly better, reaching *R*^2^ = 0.4 for *n* = 100.

We also expected some level of alignment between the asymmetry of connections. We thus repeated the same analysis, considering the *R*^2^ between (subgroup-averaged) Δ |*Ĉ*| and 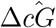. When considering *Ĉ*-MOU, The alignment increases with group size *n*, reaching values *R*^2^ ≈ 0.3 for *n >* 20 (Fig. 4c) and *R*^2^ = 0.39 for *n* = 100. The alignment between Δ |*Ĉ*| and the non-corrected 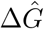 is notably inferior (*R*^2^ = 0.13 for *n* = 100). When considering *Ĉ*-rDCM, we observed a very weak alignment (Fig. 4d), remaining at the level of *R*^2^ ≈ 0.03 independently of subgroup size. In summary, the theoretically expected alignment between effective connectivity and Granger causality at the level of link strength is visible in fMRI data at a group level, and it is much stronger when EC is obtained via MOU-EC. Alignment at the level of link asymmetry was also visible at group level, but only for the corrected version of Granger causality and for MOU-EC.

Some comments about the discrepancies between *Ĉ*-MOU and *Ĉ*-rDCM are in order. At a group level, we observe good alignment between *Ĉ*-MOU and *Ĉ*-rDCM. As long as we consider subgroups of ≳ 20 participants, we observe *R*^2^ *>* 0.55 at the level of link strength and *R*^2^ *>* 0.3 at the level of link asymmetry (Fig. 4e). For the whole group of 100 participants, we obtain *R*^2^ = 0.64 at the level of link strength and *R*^2^ *>* 0.31 at the level of link asymmetry. Despite this partial convergence, the two EC methods seems to be affected by different biases. *Ĉ*-MOU estimates are biased towards 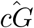, both at the level of link strength and link asymmetry. As for *Ĉ*-rDCM, we observed a strong alignment with the lagged cross-covariance *Q*^1^ = ⟨**x**(*t*)**x**(*t* + 1) ⟩ (Fig. 4f). In particular, for the whole group of *n* = 100 participants we observed *R*^2^ = 0.58 between *Ĉ*-rDCM and 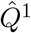; the alignment was even stronger at the level of link asymmetry (*R*^2^ = 0.85).

#### Sources and Sinks for EC and GC

Directed connectivity measures such as effective connectivity and Granger causality are often used to infer hierarchies between brain regions, based on a reconstruction of ‘sources’ (nodes with predominantly outgoing connections) and ‘sinks’ (nodes with predominantly incoming connections). We tested to what extent different measures would lead to a consistent identification of source and sink regions. To this aim, we considered the ‘net outgoing strength’ as the difference between the out-strength and the in-strength of each nodes, i.e., the total difference between outgoing and incoming link strengths. For instance, for region *i*, the net outgoing strength measured by 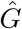 is

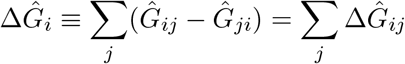

Analogous metrics can be defined for 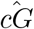, *Ĉ*-MOU and *Ĉ* − *rDCM*. For *Ĉ*, we consider two versions of this metric, one accounting for the presence of excitatory and inhibitory connections (thus using the signed link values *Ĉ*_*ij*_), and one considering the absolute link strength (hence using the absolute values |*C*_*ij*_|). These resulting maps are shown in Fig. 5a. All maps exhibit some commonalities: for example, sensorimotor regions are consistently identified as sources by all measures, while the precuneus and posterior cingulate are consistently identified as sources. However, correspondence between all maps is limited. In Fig. 5b we show the correlation (Pearson *R*) between all pairs of maps. Results are clearly in line with previous considerations. The maps obtained with rDCM taking the signed vs. absolute value are nearly equivalent (given the minor relevance of negative connections). The maps obtained with MOU are instead quite different depending on taking or not the absolute value: the maps obtained with the signed values aligns best with the rDCM map, while the map obtained with absolute values aligns with the *cG* map. Discrepancies between the maps are visible from Fig. 5c, where we show the net outgoing strength for nodes of different RSNs. While all methods find a positive net outgoing strength for SMN and DAN, and a negative net outgoing strength for CN, results for other networks strongly depend on the chosen method.

**Figure 5:**
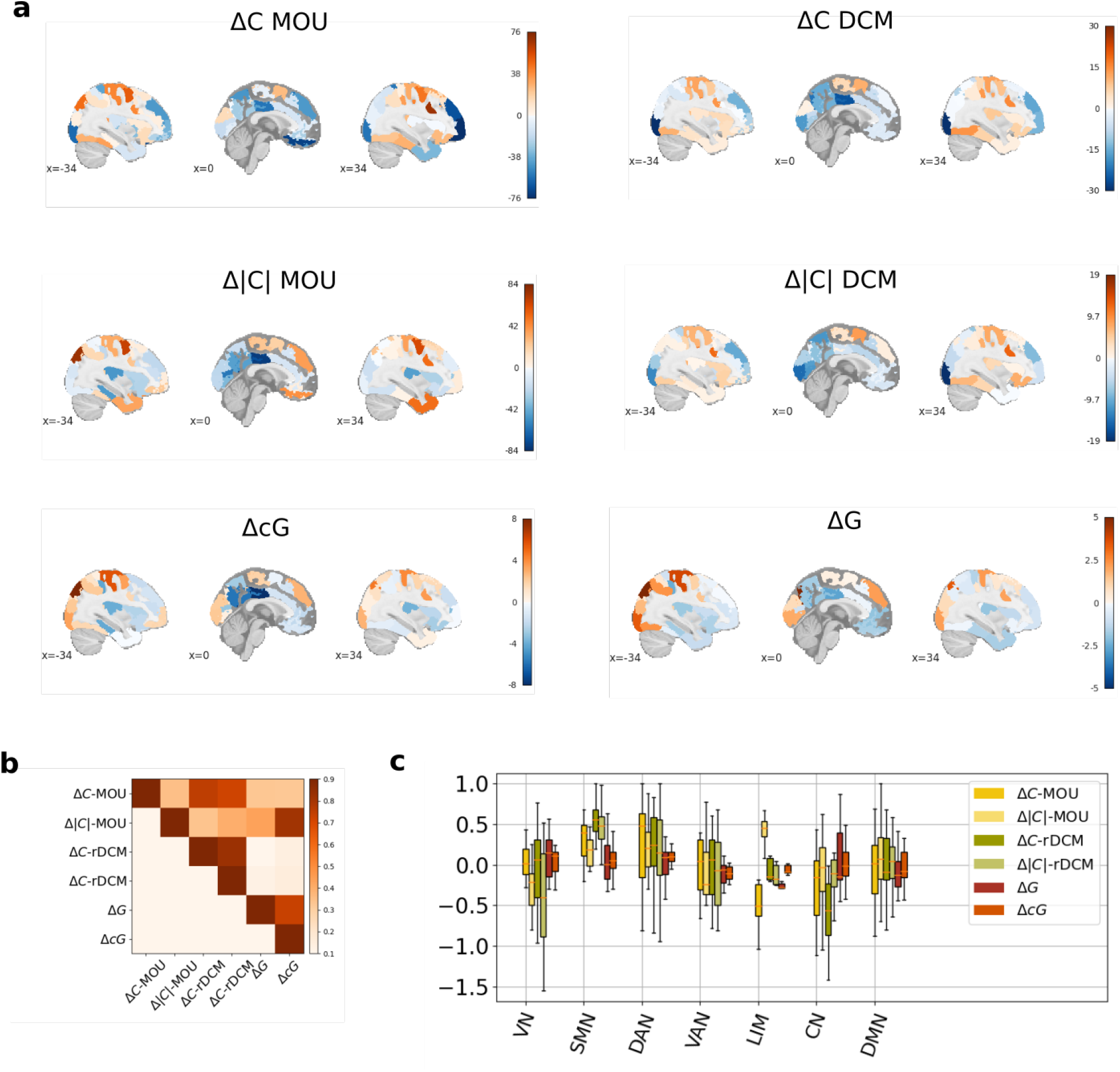
Sinks and sources for EC and GC. **(a)** Group-level maps of the difference between the out-strength and the in-strength for different metrics **(b)** Correlation between the maps **(c)** Difference between out-strength and in-strength in different resting state networks.

#### Reproducibility

We replicated the same analysis using the second fMRI session available for all subjects. In Fig. 6 we show a quantitative comparison between the results of the two sessions. In Fig. 6a we show the consistency (*R*^2^) of the estimates of *Ĉ*-MOU,*Ĉ*-rDCM, 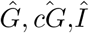 and the corresponding asymmetries at the individual level. At this level, we observed a very poor consistency (*R*^2^ *<* 0.1) between the estimates of the two sessions by *Ĉ*-MOU and 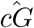. For 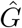 we obtained a weak *R*^2^≈0.2. Instead, for *Ĉ*-rDCM we obtained a good consistency *R*^2^ = 0.5, and an even better one for 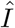 (*R*^2^ ≈ 0.6). The result for 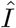 is in line with those obtained for the non-lagged and lagged covariance matrices 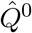 and 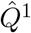. When considering link asymmetries, results of Δ*Ĉ*-MOU, 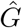 and 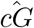 were also poorly consistent *R*^2^ *<* 0.05, while some consistency was obtained by Δ*Ĉ*-rDCM (*R*^2^ = 0.15). We hypothesize that the much larger consistency of *Ĉ*-rDCM, compared to *Ĉ*-MOU and 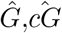 is due to the previously noticed strong convergence with *Q*^1^. In Fig. 6b we show group-level results obtained when considering subgroups of different size *n*. Averaging over at least 20 subjects yielded reliable estimates (*R*^2^ *>* 0.6) for all measures. Overall, the degree of consistency showed a hierarchy *Ĉ*-MOU, 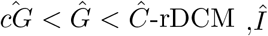. Consistency at the level of link asymmetry also obeyed the same trend, with reliable estimates (*R*^2^ *>* 0.4) starting from 20 subjects (Fig. 6C), and the same hierarchy between the different measures. When using the whole group of *n* = 100 subjects, estimates of connection strength and asymmetry were consistent at a group level (*R*^2^ *>* 0.8) for all measures.

**Figure 6:**
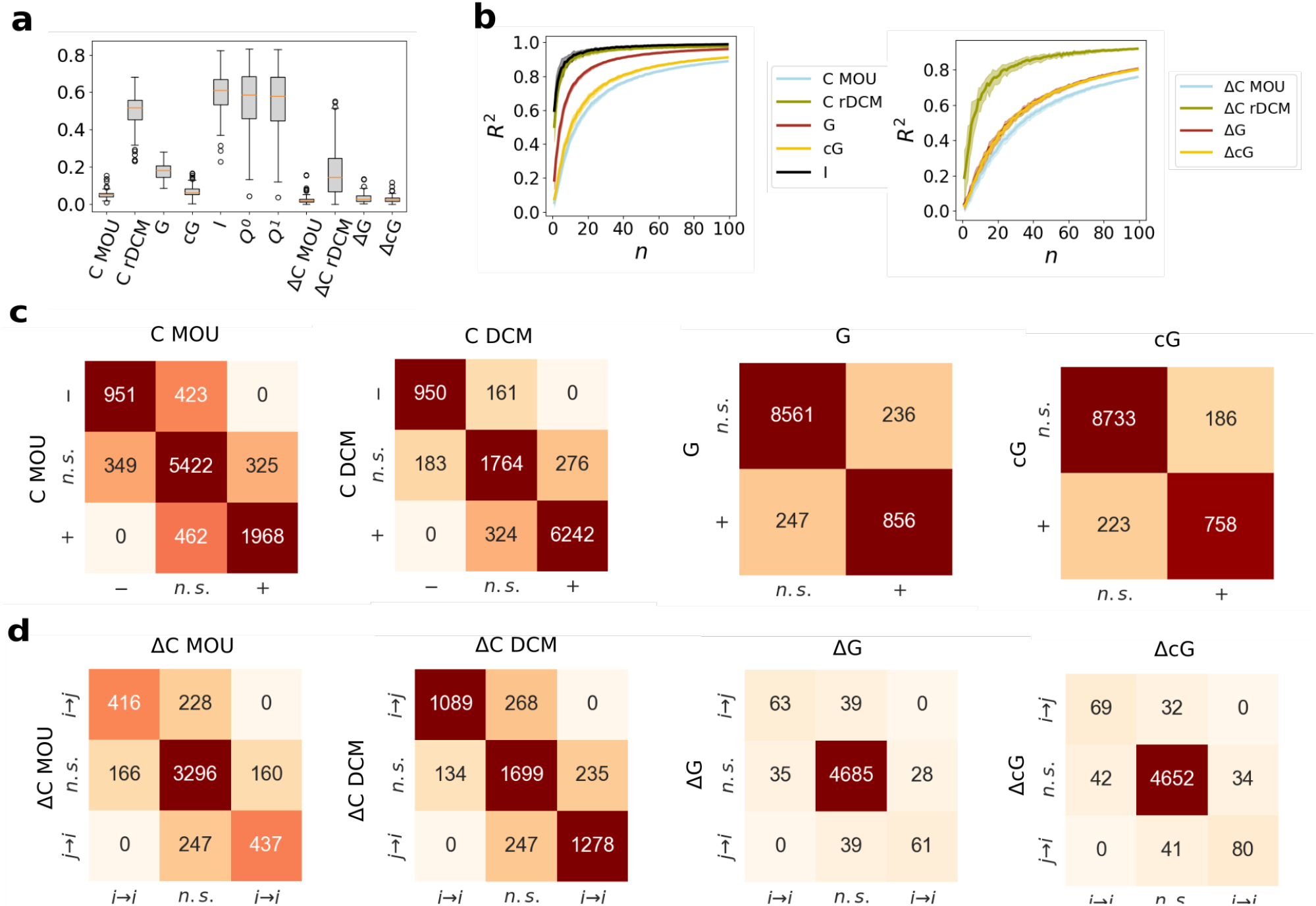
Consistency of results in two independent independent recording sessions **(A)** consistency of *single-subject* estimates across the two sessions (squared Pearson correlation). **(B)** Squared Pearson correlation *R*^2^ between group estimates of *Ĉ*, 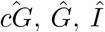 for groups of increasing size *n*. **(C)** Squared Pearson correlation *R*^2^ between group estimates of Δ*Ĉ*, 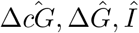 for groups of increasing size *n*. **(D)** Confusion matrix between the connection category identified in fMRI session 1 (significantly positive, significantly negative, non significant/null) and the category identified in fMRI session 2, at a group level (*n* = 100). **(E)** Confusion matrix between the connection asymmetry category identified in fMRI session 1 (significant asymmetry from *i* → *j*, significant asymmetry from *j* → *i*, non significant asymmetry) and the category identified in fMRI session 2, at a group level (*n* = 100).

We also tested for consistency in the identification of significant links. In Fig. 6c we compare the links identified as significant in the two sessions by several measures. For *Ĉ*-MOU, ≈ 80% of the links identified as significantly positive in session one was also labeled as significantly positive in session two, with the remaining 20% labeled as non-significant (and vice versa); ≈ 70% of the links identified as significantly negative in session one was also labeled as significantly negative in session two, with the remaining 30% labeled as non-significant (and vice versa); no link was labeled as significantly positive in one session and significantly negative in the other. An even more consistent picture was obtained for rDCM: ≈ 95% of the links identified as significantly positive in session one was also labeled as significantly positive in session two, with the remaining 5% labeled as non-significant (and vice versa); ≈ 90% of the links identified as significantly negative in session one was also labeled as significantly negative in session two, with the remaining 10% labeled as non-significant (and vice versa); no link was labeled as significantly positive in one session and significantly negative in the other. For 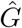 and 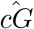, ≈ 80% of the links identified as significantly positive in session one was also labeled as significantly positive in session two.

Finally, we tested the consistency in the identification of links with significant asymmetry (Fig. 6d). For *Ĉ*-MOU, ≈ 65% of the links displaying a significant link asymmetry in session 1 also presented a significant asymmetry in the same direction in session 2, with the remaining 35% displaying a non-significant asymmetry (and vice versa). For *Ĉ*-rDCM, ≈ 85% of the links displaying a significant link asymmetry in session 1 also presented a significant asymmetry in the same direction in session 2, with the remaining 15% displaying a non-significant asymmetry (and vice versa). For 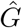 and 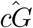, approximately 65% of the links displaying a significant link asymmetry in session 1 also presented a significant asymmetry in the same direction in session 2, with the remaining 35% displaying a non-significant asymmetry (and vice versa). No measure identified asymmetries in opposite directions in session 1 and 2.

In summary, results are highly replicable, but only at a group level. Overall, rDCM produced more consistent results than MOU-EC and Granger causality.

## Discussion

A fundamental objective of system and network neuroscience is to uncover the graph of causal inter-connections between areas of the human brain. Substantial information about directional connections in the brain can be learned from time-lagged relationships between brain areas observed in fMRI [17]. Traditional, non-directional functional connectivity cannot characterize asymmetries between reciprocal connections, which underlie a key principle of brain organization, hierarchical processing [57, 58]. Directional connectivity methods overcome these limitations, elucidating the task-related modulation of functional interactions [59], hierarchical relations underlying cognitive functions (such as working memory [60, 61], cognitive control [62] and language [43]), and pathological alteration of functional hierarchies [28, 63] - leading to improved biomarkers for clinical classification [42].

Despite the availability of several methods to analyze directional connectivity [64], the majority of studies employ effective connectivity (EC) or Granger causality (GC). The state-of-the-art wisdom about these two methods is somewhat contradictory. On the one hand, the methodological literature has often stressed their differences [32], insisting that they rest on different premises: EC aims to retrieve a latent dynamical model, while GC is based on statistical dependencies. On the other hand, in applications the two techniques are used quite interchangeably, EC or GC links being generally interpreted as directional couplings between brain areas, with an implicit intuition that large effective connections should be reflected into large values of Granger causality and vice versa.

### Theoretical relations explaining similarities and differences between GC and EC

The first contribution of this work is to provide a clear theoretical framework to understand similarities and differences between EC and GC in the context of brain network reconstruction in fMRI. Common EC approaches for whole-brain fMRI assume a linear, stochastic dynamical model driven by Gaussian noise, i.e., a multivariate Ornstein-Uhlenbeck process. Similarly, GC approaches assume a linear multivariate autoregressive (MVAR) process with Gaussian noise. Assuming the MVAR to be first-order (as commonly done in fMRI studies [20]), we leveraged an explicit mapping between the continuous-time MOU and the discrete time MVAR to derive analytical relations between EC and GC (Eqs.(32),(33) and Fig. 1), valid between *model* values of the two quantities (which of course differ from *estimates* obtained from finite samples). These relations allowed us to formulate clear predictions about the EC-GC correspondence.

Firstly, the implication large EC ⇒ large GC critically depends on the relation between two time scales: the sampling time (Δ*t*; interval between two successive observations, corresponding to TR in fMRI) and the process time (*τ*; the time scale of the stochastic ODE system underlying EC analysis, related to the autocorrelation time of the dynamical system). When the process is fast (*τ* ≪ Δ*t*), the values of the (discrete) observed time series at the previous time point are poorly predictive of the next time point. Thus, regardless of the strength of effective connections, GC remains low (as underscored by previous work [55]). The presence of a strong effective connection reflects in an “instantaneous” correlation that cannot be predicted on the basis of previous time points, which is captured by the “instantaneous (Granger) causality” (IC), a non-directional connectivity measure that is often neglected but is part of the original Granger-Geweke formalism [47]. When the process is slow (*τ* ≫ Δ*t*), the values of the time series at the previous time point are strongly predictive of the next time point. In this case, the presence of a large effective connection reflects in large values of GC (while IC is very small). In fact, the analytical relations (Eqs.(32),(33)) show that both GC and IC are (roughly) proportional to the square of EC, but for *τ* ≪ Δ*t* the proportionality constant is much stronger for IC than GC (and vice versa for *τ* ≫ Δ*t*). Thus, our study shows that large effective connections are reflected in large values of Granger causality *or* instantaneous causality.

Secondly, a direct relation between EC and GC link strengths holds only if all areas have equal signal variance, corresponding to a homogeneous level of random input. Otherwise, the relation is modulated by the ratio of the input noise affecting the two endpoint nodes. As an example, for symmetric effective connections between two areas, the GC from the area with higher variance will be larger than in the reverse direction. This is not necessarily a pitfall of GC, that was conceived as a measure of “influence”: in presence of equal connections, the influence of an area with larger variance (signal power) is stronger. However, it implies that GC cannot be considered a measure of coupling, like EC. In presence of non-homogeneous signal power, it is still possible to recover a quadratic relation between EC and GC/IC (Eqs.(34),(35)) by “correcting” GC estimates by simple factors accounting for the heterogeneity in nodal input.

Thirdly, the EC-GC relation is non-monotonic (approximately quadratic, Eqs.(32),(33)). Effective connections can be positive or negative, and larger values of GC correspond to stronger values of EC in *modulus* (i.e, strongly positive or strongly negative). This is again consistent with the interpretation of GC as a measure of influence: strongly “excitatory” or strongly “inhibitory” effective connections both determine a large causal effect from the source node to the target node. Correspondingly, EC-GC alignment at the level of link asymmetry can be observed only if the definition of the EC asymmetry involves the modulus of the connection strength (Fig. 1), identifying the direction of the strongest connection (whether positive or negative).

### Effects of sampling

The second contribution of this work is to give predictions about the level of the EC-GC alignment that one may expect in finitely sampled time series. The analytical relations obtained hold at the theoretical level, i.e., between model couplings of a continuous-time MOU process and model GC values of the corresponding discrete-time MVAR process. In practice, EC and GC *estimates* obtained from finite samples differ from the model values.

With the aid of numerical simulations of artificial networks, we discussed to what extent the theoretical EC/GC relations could be observed from finite-length data, depending on the sampling rate and the sampling length (Fig. 2). For estimation, we used the two most common methods for whole-brain EC estimation (MOU-EC and regression DCM) and a popular approach for GC estimation (the covariance-based approach). The expected relations were clearly observed only for a sufficiently fast sampling rate that matches the process time scale (Δ*t* ≤ *τ*). Moreover, a sampling length of ℒ = 10^4^ was required, which amounts to 10 times the length of a typical recording session. For ℒ = 10^3^ (the typical sampling time of an fMRI session), we observed a strong EC/GC correlation only when using MOU-EC, but not rDCM. This is probably due to common estimation biases between covariance-based GC and MOU-EC, which both depend on empirical covariance matrices. This analysis shows that GC-EC correlation can emerge only in unusually long recordings (such as [65]) or, more plausibly, at the group level (neglecting inter-individual variability). Thus, we predicted that group-level analyses using at least 10 subjects (order of magnitude) should be considered to observe the EC-GC alignment in fMRI data. To observe alignment at the level of connection asymmetry, this requirement becomes even stricter: at least *N* ~ 100 subjects should be used to reach the theoretically predicted level of correlation (*R*^2^ ≈ 0.4).

### EC/GC comparison in fMRI data

The third contribution of this work is an assessment of the consistency of EC and GC in fMRI data. Although some methodological studies restricted attention to simulated network activity to compare directional connectivity methods [55, 64], using real data is crucial, as certain assumptions behind synthetic data (e.g. Gaussianity of generated signals) might only be loosely respected, if not entirely violated.

In agreement with predictions from simulated data, we observed EC-GC correlation *at the group level* (Fig. 4) when pooling at least ~15 subjects together. Consistently with the simulation results, the GC-EC relation was much stronger for MOU-EC than rDCM (*R*^2^ *>* 0.6 vs *R*^2^ *>* 0.25). Overall, EC analysis detected a much larger number of significant connections at a group level (38% for mou-EC, 77% for rDCM, compared to 9% for GC; fig. 3). The overwhelming majority of significant GC links were identified as significant by EC (both my MOU-EC and rDCM), while the converse did not hold. This demonstrates that GC is more conservative than EC in identifying significant connections at a group-level. At the level of link asymmetry, consistency between the two methods was achieved only for MOU-EC, and only if a group of 20 subjects or more was employed (fig. 4). Congruently, when aggregating asymmetry at the nodal level (by computing the difference of the out-strength and in-strength of each node), we obtained a good alignment between GC and mou-EC (*R*^2^ ~ 0.6), but not between EC and rDCM-EC (*R*^2^ ≈ 0); Fig. 5). This implies that GC and EC could give widely different pictures of functional hierarchies, depending on the method chosen for EC estimation.

Our test-retest analysis (Fig. 6 also allowed us to assess the reliability of single-subject and group estimates of EC and GC. GC estimates were not reliable at the single-subject level, but we obtained a good test-retest consistency *R*^2^ = 0.8 at the group level for groups of at least ~ 20 subjects. The reliability of EC estimates depended strongly on the chosen estimation method. RDCM yielded relatively consistent estimates (*R*^2^ = 0.5) even at the individual level and well consistent estimates (*R*^2^ *>* 0.8) for groups of at least 5 subjects, whereas MOU-EC was poorly consistent individually (*R*^2^ *<* 0.1) and achieved good group consistency (*R*^2^ *<* 0.8) only for groups of at least 50 subjects. This striking difference should be attributed to the different estimation procedures of the two methods. In the absence of a ground truth, we hesitate to conclude that rDCM yields “better” estimates. In fact, we observed that rDCM estimates correlated very strongly with the lagged covariance matrix (*R*^2^ ~ 0.6). This strong bias may improve the test-retest consistency, but also impoverish the meaningfulness of rDCM EC, which appears to be barely different from a time-lagged FC. We stress that the lack of reliability for MOU-EC concerns individual connection estimates; this does not necessarily contradict previous work that leverages information from the whole network of connections to obtain robust prediction of clinical condition [40–42].

### Future work and potential limitations

In this study, we did not investigate or model hemodynamics. The effect of hemodynamics could be multiple: it could introduce deviations from the model at the level of observed BOLD time series (such that a MOU or first order MVAR no longer provide an accurate description of the data), or bias timing relationships due to regional differences in the hemodynamic response, hence biasing EC/GC estimates (in particular estimates of connection asymmetry). In principle, one should estimate EC and GC from “neural” time series obtained after a deconvolution of the hemodynamic response. Additional work is needed to appreciate the effect of the hemodynamic response on the GC/EC relation, e.g. applying blind approaches to deconvolve the hemodynamic response from resting-state fMRI data (such as [66, 67]). Another possible limitation of this study is the assumption of a diagonal noise covariance matrix in the MVAR model: in the presence of a large common input between regions, the noise should be modeled as having non-diagonal covariance ∑, which could reflect into larger values of the instantaneous Granger causality *I*. This could explain the anomalous (large) IC values observed in fMRI data, despite the fact that *τ* ~ Δ*t*. Reliably inferring a non-diagonal noise covariance poses additional challenges [68, 69]. Future work will better investigate the relation between *C*, ∑ and *G, I* in the case of correlated noise.

## Conclusion

Our study addressed the consistency of GC and EC in the reconstruction and analysis of directional brain networks from neuroimaging time series. In the context of the first-order autoregressive models typically used in fMRI, GC and EC share common assumptions and are mathematically related. The relation is non-trivial due to the presence of negative effective connections and unequal noise variance on different network nodes. Moreover, the relation can emerge only in the limit of large sampling - which, due to practical limitations in fMRI acquisition, means that in practice it can be observed only at the group level for at least 10 − 20 subjects. Furthermore, the GC-EC alignment is strongly affected by the method used for EC estimation (MOU-EC giving much stronger alignment than rDCM). Taking these factors into account in the analysis of real fMRI recordings, we showed that the two methods yield a relatively consistent description of directional brain networks at a group level.

## Data and Code Availability

All simulated data and code used in the present manuscript is available at https://github.com/micheleallegra/GC-and-MOU. The HCP data are part of the 1200 Subjects Release (S1200) and are publicly available on the ConnectomeDB database https://db.humanconnectome.org.

## Acknowledgments

M.A. and A.B. were supported by FLAG-ERA JTC 2017, grant ANR-17-HBPR-0001, “Brainsynch-Hit”. MA was supported by the Italian Ministry for University, through the 2022 PRIN program, project 2022HSKLK9, “Unveiling the role of low dimensional activity manifolds in biological and artificial neural networks”. MG received support from the French government under the France 2030 investment plan, under the Agence Nationale de la Recherche grant (ANR-22-CPJ2-0020-01) and as part of the Initiative d’Excellence d’Aix-Marseille Université – A*MIDEX (AMX-22-CPJ-01). MG was also partly supported by the European Union Horizon 2020 Grant No. 945539 (Human Brain Project SGA3). AB was supported by the Agence National de la Recherche, Grant ANR-18-CE28-0016 and by the European Union’s Horizon 2020 Framework Programme for Research and Innovation under the Specific Grant Agreement No. 945539 (Human Brain Project SGA3).

## Supporting information - Appendix

### Multivariate autoregressive (MVAR) models and Granger Causality

A 1^st^ order *N*-dimensional multivariate autoregressive (MVAR) process is defined for discrete time *t* (with steps Δ*t*) by

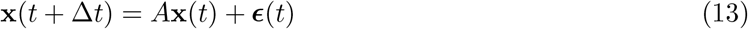

where ***ϵ***(*t*) is Gaussian noise with zero mean and covariance matrix *S*,

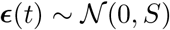

and ||*A*|| ≤ 1 for stability. The innovation, or noisy input, *ϵ*_*j*_(*t*) in Eq. (2) corresponds to the residual of the linear regression

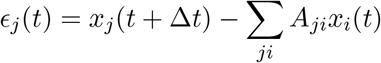

The variance of *ϵ*_*j*_(*t*) (corresponding to a diagonal element of matrix *S*) reflects the “magnitude” of the residual, which measures how well past values of the time series can predict the next value at *j, x*_*j*_(*t* + Δ*t*), namely

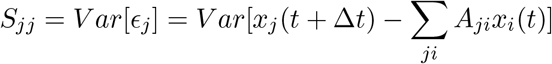

To quantify the Granger causal effect of node *i* on node *j*, one can measure the relevance of *x*_*i*_(*t*) in predicting *x*_*j*_(*t* + Δ*t*). To this aim, one defines a a “reduced” MVAR process where the influence of node *i* is removed:

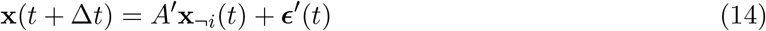

where **x**_¬*i*_(*t*) is obtained from **x**(*t*) by removing its *i*-th component and ***ϵ***′(*t*) is Gaussian with zero mean and covariance matrix *S*′,

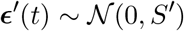

Again, the magnitude of the residual 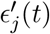 can be assessed via its variance

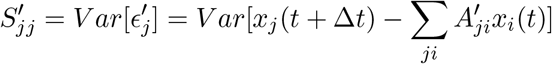

The effect of node *i* on node *j* defined by *Granger causality* (GC) [23, 47] is given by the log-ratio of the variances

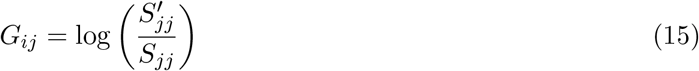

Note that we consider the conditional version of GC, meaning that the linear regression in Eq. (3) includes all remaining nodes in the network. Also note that Eq. (4) defines a “model” GC, which differs from its *estimates* obtained from finite data and noisy covariance matrices (see below). We can show an approximate relation between the Granger Causality *G*_*ij*_ and the MVAR coefficient *a*_*ji*_. Assuming that *A*′ ≃ *A*^¬*i*^ (where *A*^¬*i*^ is simply *A* without the *i*-th column), one obtains

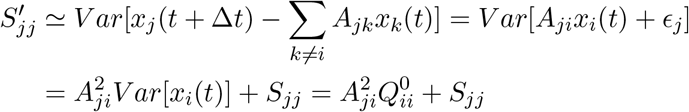

using the further assumption of the statistical independence between *x*_*i*_(*t*) and *ϵ*_*j*_. Therefore we have

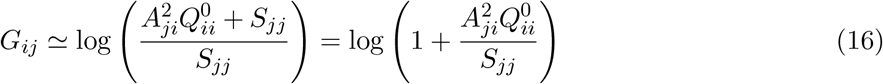

hence *G*_*ij*_ is approximately a monotonic function of 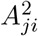. For sufficiently small 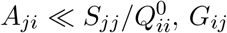, *G*_*ij*_ is approximately a quadratic function of 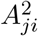.

In addition to the standard Granger causality (4), Geweke [47] defined the “instantaneous” Granger causality *I*_*ij*_, which compares the magnitude of the innovations when considered jointly or separately. The innovations {*ϵ*_*i*_(*t*), *ϵ*_*j*_(*t*)} jointly have a covariance matrix

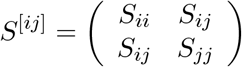

The total magnitude of the joint innovations can be measured as log(det *S*^[*ij*]^). If one considers the innovations independently, one obtains instead log *S*_*ii*_ + log *S*_*jj*_. The instantaneous Granger causality is defined as

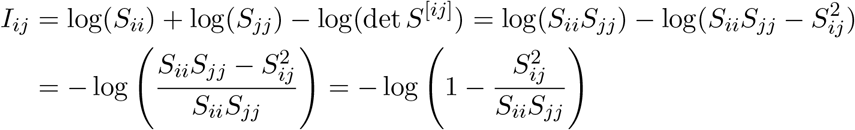

While the instantaneous causality is often discarded in Granger causality analyses, it may capture a large part of the interdependence between two time series.

### Covariance-based GC

A standard method to estimate *G*_*ij*_ from the data is by inferring the parameters of the MVAR (2) and the reduced MVAR (3). An alternative and computationally simple approach exploits a relation between the variance of the MVAR residuals and covariance terms, assuming Gaussian innovations [53, 70, 71]. Indeed, it can be shown that

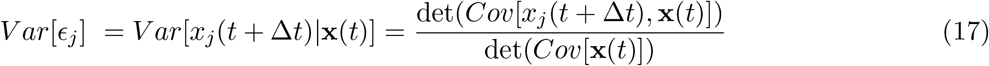

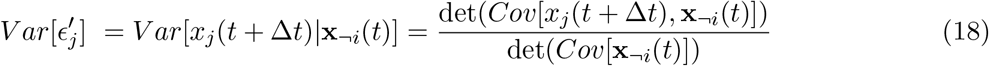

Furthermore, we have:

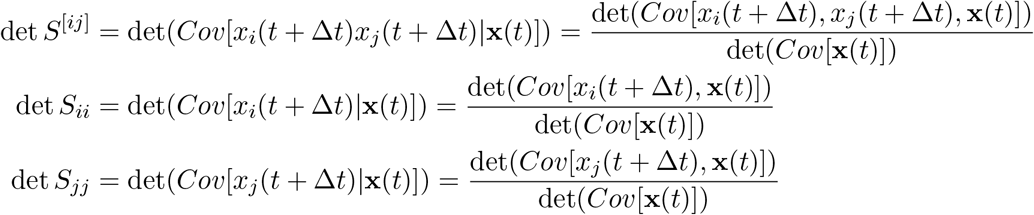

Thus, exploiting covariance relations, we can express *G*_*ij*_ and *I*_*ij*_ as follows:

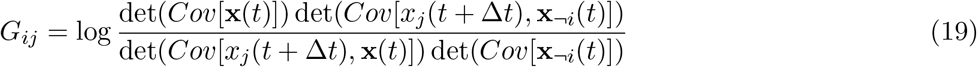

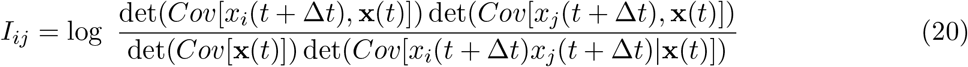

Thus, *G*_*ij*_ and *I*_*ij*_ can be both expressed in terms of elements of the 0-lagged and the Δ*t*-lagged covariance matrices

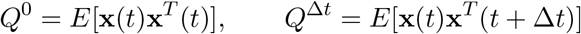

Note that for Gaussian systems, there is a relation between the entropy and the covariance matrix,

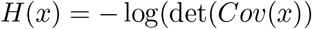

Using this relation, one can show that the covariance-based GC is equivalent to the transfer entropy [22].Granger causality measures can therefore be formulated in completely information-theoretical terms, based on entropy estimates [28, 53].

Estimates 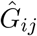 and 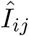 can be obtained from finite-sampling estimates of the covariance matrix,

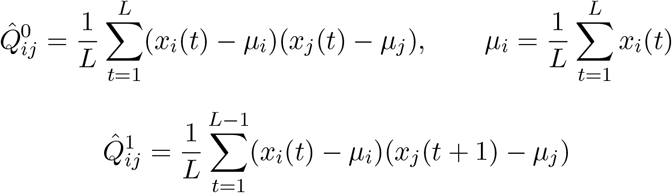

Before computing covariance-matrices, we applied the cop-norm transformation on the signal of each region, 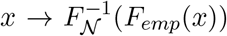 where *F*_*emp*_(*x*) is the empical cumulative distribution function of *x*, and *F*_𝒩_ (*x*) is the cumulative distribution function of a standard normal. This transformation corrects for possible non-Gaussianity of the signal distributions, assuming a Gaussian copula [72].

### Multivariate Ornstein-Uhlenbeck (MOU) process and relation to MVAR

Another approach to characterize the causal effect of node *i* on node *j* is to assume a generative dynamical model in continuous time, and assess the value of the *i, j* coupling that can be seen as *model effective connectivity*. Gilson et al. rely on the multivariate Ornstein-Uhlenbeck process (MOU) given by

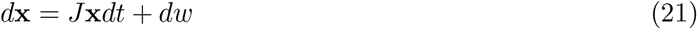

Where

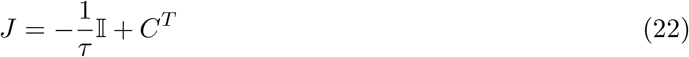

with the process time constant *τ >* 0 and the identity matrix 𝕀. Here *w* is a Wiener process (akin to white noise) corresponding to a diagonal covariance matrix ∑ with input variances 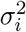 and zero input cross-covariance, such that 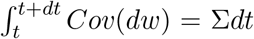. At equilibrium, the covariance matrix *Q*^0^ for **x**(*t*) following Eq. (1) satisfies the (continuous) Lyapunov equation

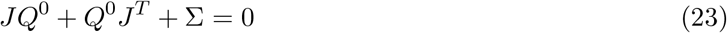

Furthermore, the 0-lagged and Δ*t*-lagged covariance matrix *Q*^0^ and *Q*^Δ*t*^ obey the relation

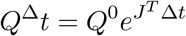

with Δ*t* a given time lag. Therefore, one can use the matrix logarithm to obtain

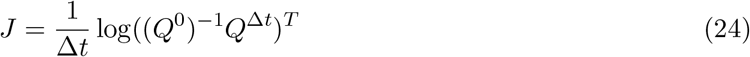

The “causal” effect of node *i* on node *j* is quantified here by the *model effective connectivity C*_*ji*_. In practice, the matrices *Q*^0^ and *Q*^Δ^*t* in Eq. (24) can be replaced by their empirical counterparts 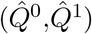 calculated from the data to obtain estimates 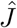, *Ĉ* of hence *J, C*. In this work, we rely on a more elaborate estimation procedure based on a gradient descent (or Lyapunov optimization) to robustly estimate *C* for fMRI data that typically consist of a limited number of time points due to the sampling rate [36]. In essence, it uses a partial differentiation of Eq. (24) to iteratively optimize 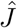 and 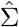. Here an important point to note is that the matrix logarithm can yield complex values for 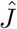, while the iterative optimization keeps the matrix elements real-valued.

For any given MOU process (1), we can build an equivalent 1^st^ order MVAR process (2). Indeed, by integrating Eq. (1) for a given lag Δ*t*, one obtains

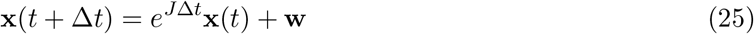

where 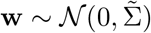 is a Gaussian noise with covariance matrix

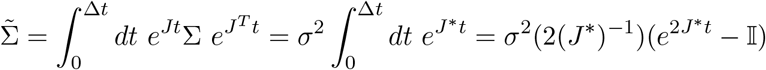

where 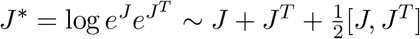.

However, for a given 1^st^ order MVAR process, the equivalent MOU process corresponds to

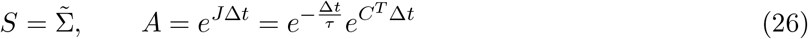

which implies the following constraints on *S* and *A*:

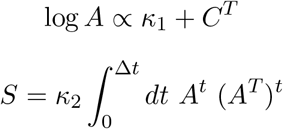

where *κ*_1_ *<* 0, *κ*_2_ *>* 0 are constants, and *C* is a real matrix with null diagonal, *C*_*ii*_ = 0.

In sum, any MOU process (1) with real coefficients is associated with an equivalent MVAR process (2) with real coefficients. The reciprocal, however, is not true, as an arbitrary matrix *A* will not yield in general a real-valued, but a complex-valued matrix *J* = log(*A*) because of the matrix logarithm. We will refer to a *MOU-compatible MVAR* when the process can be associated to a real-valued *J*.

### Theoretical relation between GC and EC

Assuming compatibility, the multivariate dynamics underlying the estimation of GC and EC are fully consistent. We can therefore compare *C*_*ij*_ and *G*_*ij*_, which both measure the effect of node *i* on node *j*. The following analysis relies on the approximation of the matrix exponential by a first-order expansion:

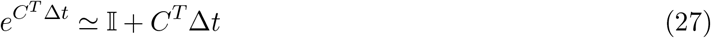

Eq. (27) holds under the following condition

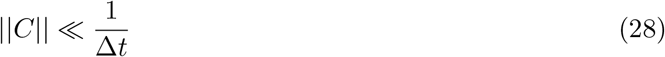

where || · || denotes matrix 2-norm. Note that this condition can be formulated considering the dominant eigenvalue of *C*: 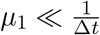 with *µ*_*i*_ being the real parts of the eigenvalues of *C* in decreasing order (*µ*_1_ ≥ *µ*_2_ · · · ≥ *µ*_*N*_). Also note that the eigenvalues of *J* are equal to those of *C* shifted by 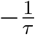, so the stability of the resulting network dynamics requires that the real parts of *µ*_*i*_ are smaller than 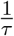 (again, it is sufficient to check only the dominant eigenvalue). We thus distinguish three regimes,

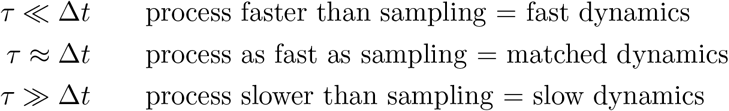

In the slow case, Eq. (28) is always satisfied since 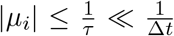. In the matched case, Eq. (28) is satisfied whenever

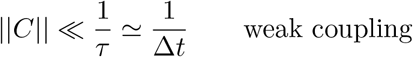

becoming more constraining in the fast case

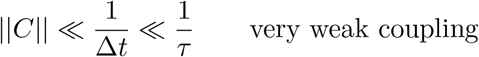

In practice, condition (28) can often be considered to be satisfied in both the slow and the matched cases, essentially because large values of *C*_*ij*_ are incompatible with stability, so ||*C*|| ≪ 1*/τ*. However, fast dynamics might break the assumption.

We can show that, provided Eq. (27) holds, there exist quadratic relations between IC, GC and EC. As detailed in Eqs. (29) and (31) in S1Text.

If Eq.(27) holds, we have quadratic relations between IC, GC and EC. Indeed, 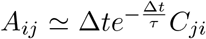 and

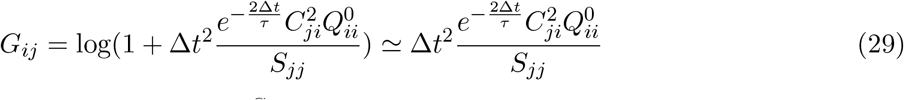

Under the same approximation we also have *e*^*Ct*^ ≃ 𝕀 + *Ct* for *t* ≤ Δ*t* so

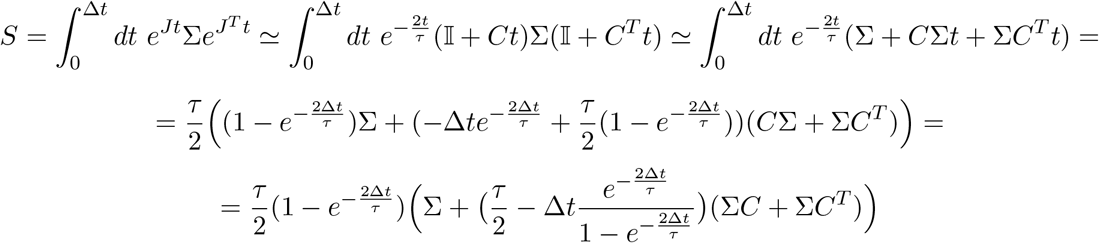

Note that 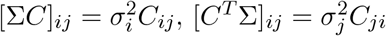. Hence,

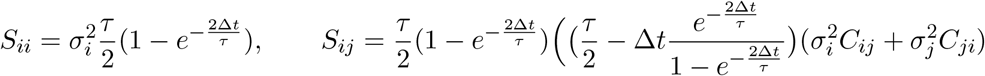

Furthermore, from Lyapunov equation one gets

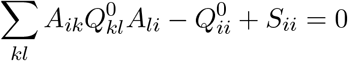

and to 1st order 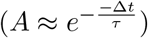

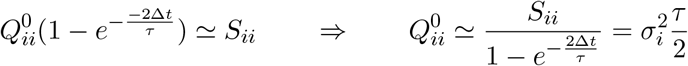

which yields, using the previous expression of *S*_*ii*_,

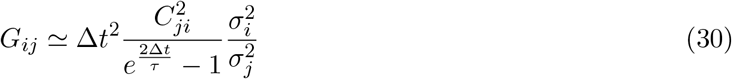

For the instantaneous causality, one obtains

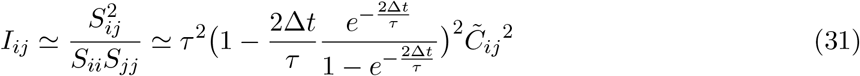

with 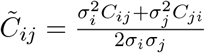.

, we have

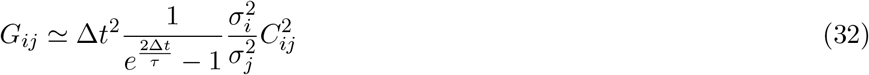

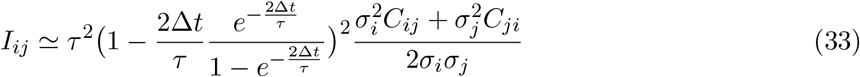

with 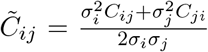 being a mean of the reciprocal connection weights between *i* and *j* weighted by the input variances. If the input noise is homogeneous across all nodes, *σ*_*i*_ = *σ*, these relations further simplify:

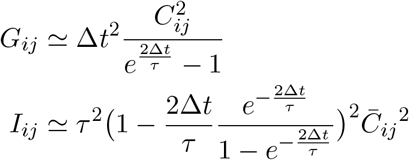

with 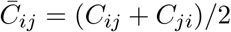 being the symmetrized effective connectivity between *i* and *j*. So, assuming equal input noise on all nodes and neglecting asymmetries, there is an approximately quadratic relation between the symmetrized effective connectivity *C*_*ij*_ and *I*_*ij*_. In particular, these relations mean that Granger causality provides an estimate of the quadratic interaction between nodes governed by continuous dynamics (i.e., having a MOU as a generative process). In the general case of inhomogeneous noise variance, we can nevertheless retrieve the approximately quadratic relation using a “corrected” versions of *G* and *I*:

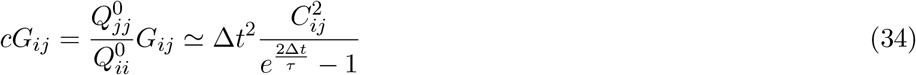

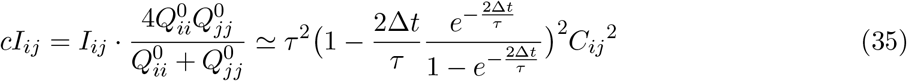

Here we have used the fact that the node variance 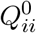 is strongly related to the corresponding input variance 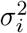. This holds in practice for weak coupling and can be seen via the Lyapunov equation (23) where *J* is then dominated by its diagonal elements 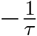, yielding 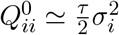.

A last quantity of interest is the ratio between Granger causality and instantaneous causality:

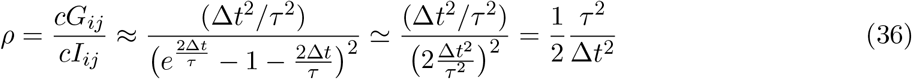

This means that for slow dynamics we have *cG* ≫ *cI*, whereas *cG* ≪ *cI* for fast dynamics (note that this also true for the uncorrected versions *G* and *I*). Thus, *G* and *I* do not always reflect the underlying network connectivity. In other words, when a continuous MOU model is a valid generative model for the observed time series, the information about *C* is differentially captured by *G* and *I*, crucially depending on the dynamical regime.

## Supplementary figures

**Figure S1:**
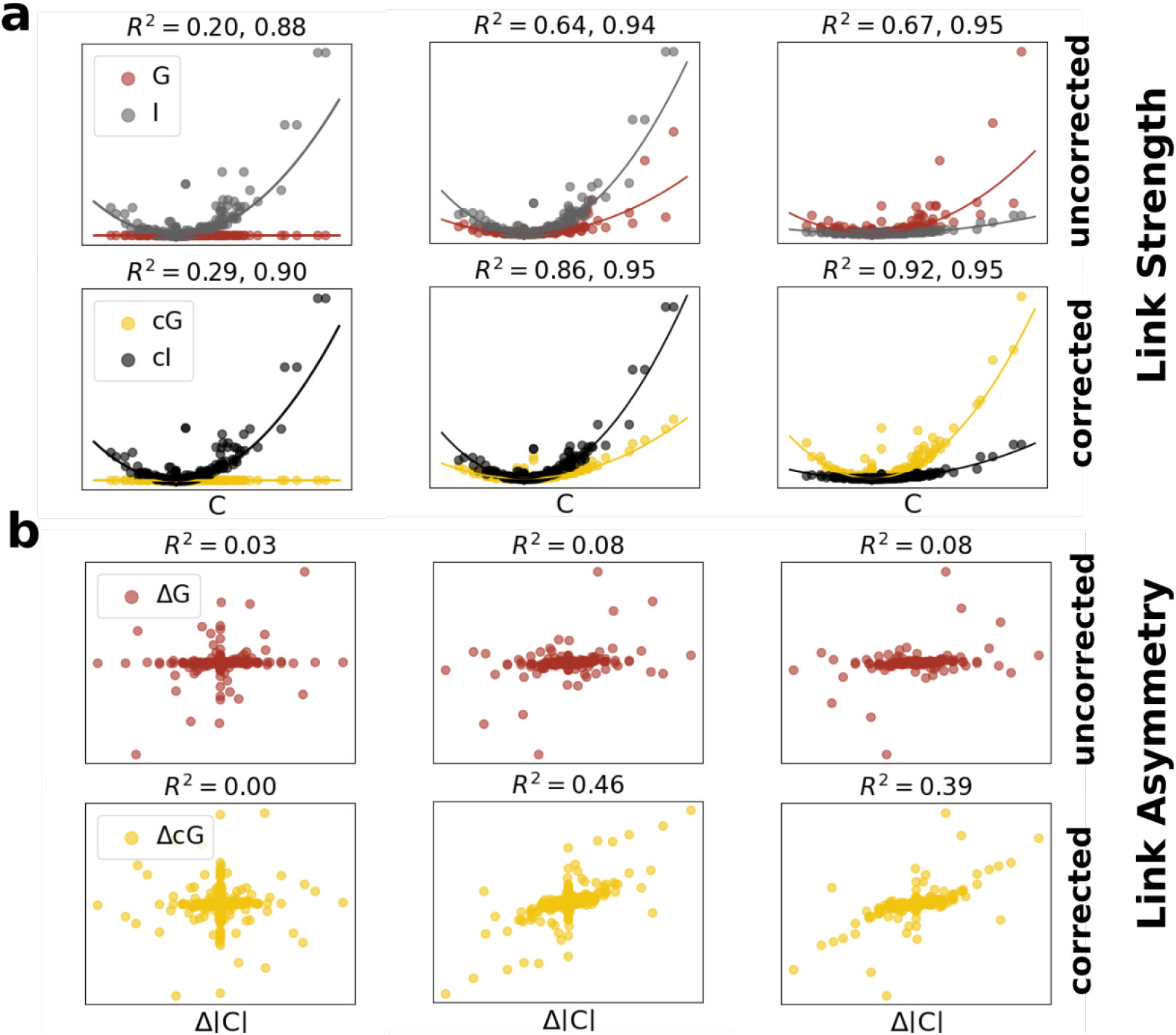
Theoretical relations between model EC and non-conditional GC/IC. We considered a random network of *N* = 40 nodes evolving according to the MOU of *τ* = 0.1, 1, 10. In **(A-C)** we show the relation between the EC weights and the corresponding values of GC and IC. The GC and IC were obtained from the ideal covariance matrices 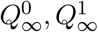 obtained in the limit of infinite observation length, *T* → ∞. Each dot corresponds to a pair (*i, j*), and straight lines to the approximate quadratic scalings, 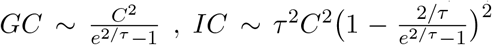. For all valued of *τ*, the approximate quadratic relation are well satisfied. The relative importance of IC vs. GC depends on the value of *τ*, with IC prevailing at low *τ* and GC at large *τ*.

**Figure S2:**
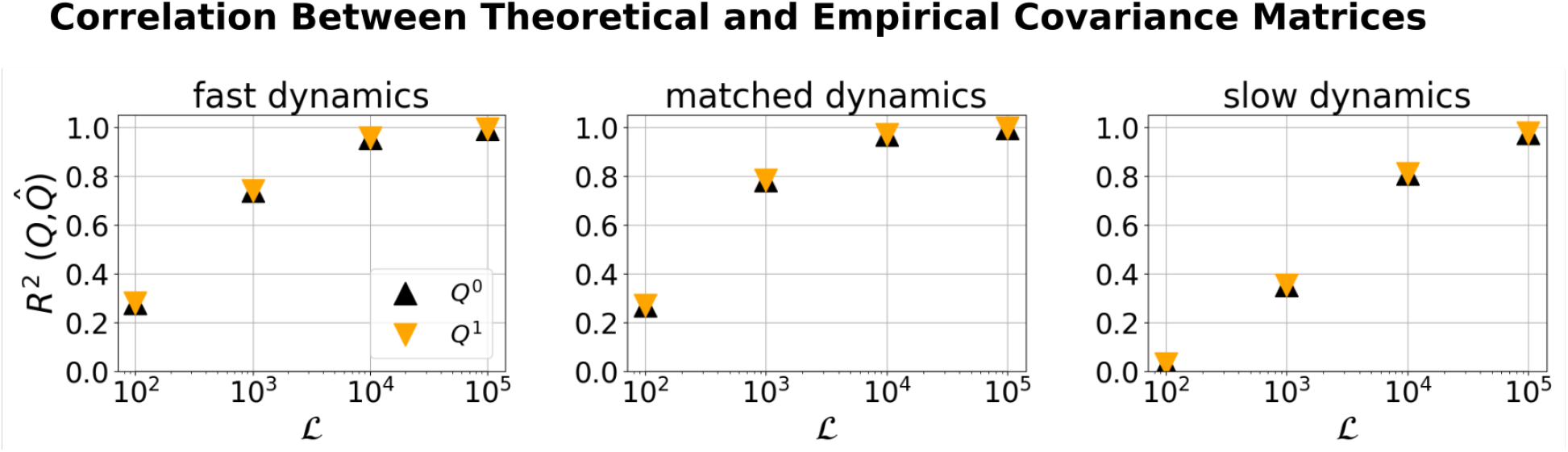
Error in covariance matrices for finite sampling time. We considered the same random network as in Fig. 2. For finite sampling length ℒ *<* ∞, the covariance matrices *Q*^0^, *Q*^1^ are affected by an estimation error, whose magnitude decreases with ℒ. We show here the relation between the ideal *Q*^0^, *Q*^1^ and the empirical 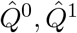, for different values of *τ* and ℒ, measured in terms of squared Pearson correlation *R*^2^.

**Figure S3:**
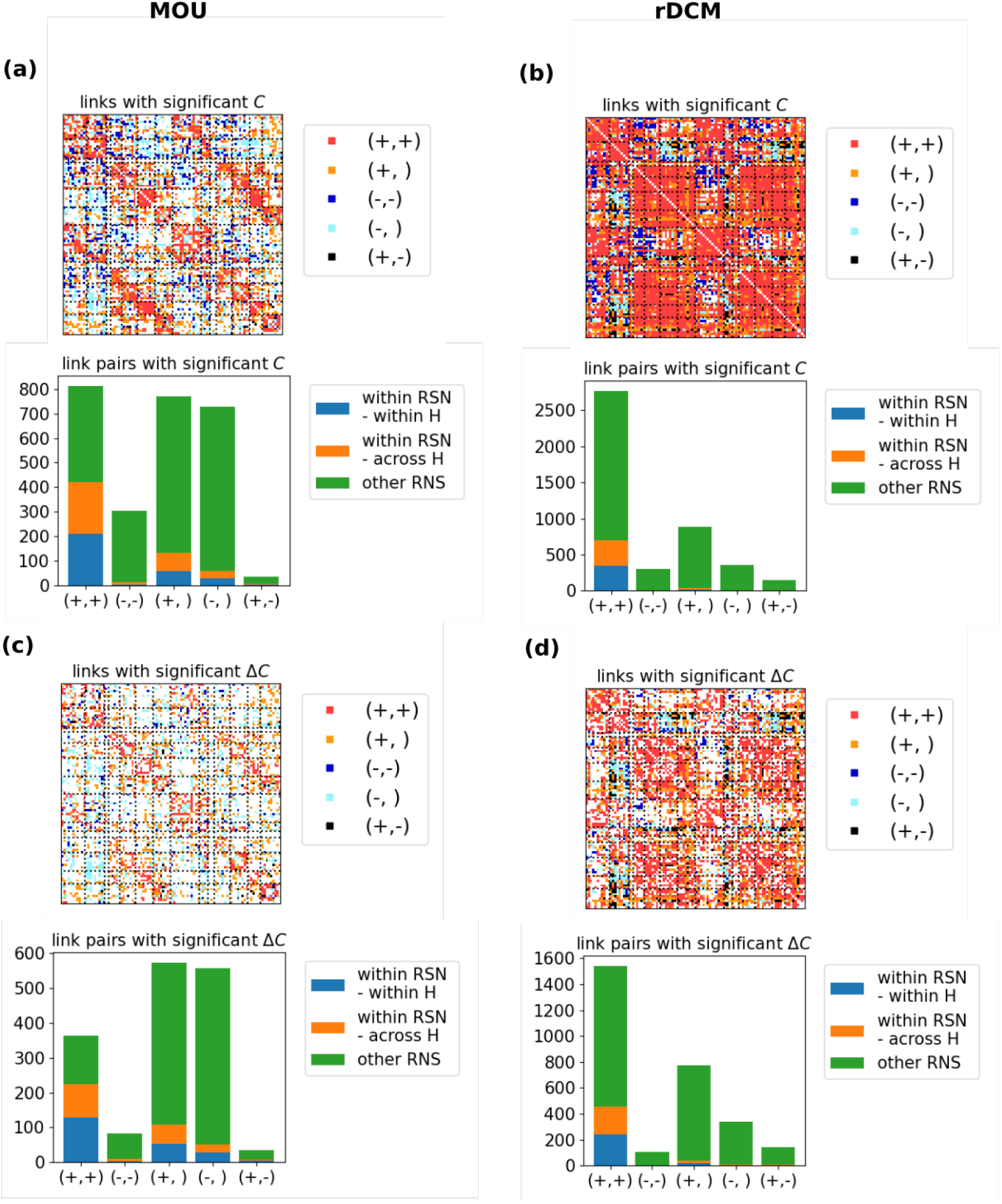
Effective Connection sign and asymmetries. We considered all pairs of reciprocal connections, and divided them into *sign categories* according to their sign: (+, +) both connections are significantly positive (+,) one connection is significantly positive and the reciprocal is non-significant (−), both connections are significantly negative (−,) one connection is significantly negative and the reciprocal is non-significant (+, −) one connection is significantly positive and the reciprocal is significantly negative. Furthermore, connections pairs were divided in *network categories* depending on the connected areas: i) areas belonging to the same RSN and the same hemisphere ii) areas belonging to the same RSN but different hemispheres iii) areas belonging to different RSNs. **(A)** We show all significant link pairs, with color depending on their sign category **(B)** For each sign category, we show the fraction of significant connections belonging to each network category. **(C)** We show all link pairs with significant asymmetry, with color depending on their sign category **(D)** For each sign category, we show the fraction of connections with significant asymmetry belonging to each network category.

**Figure S4:**
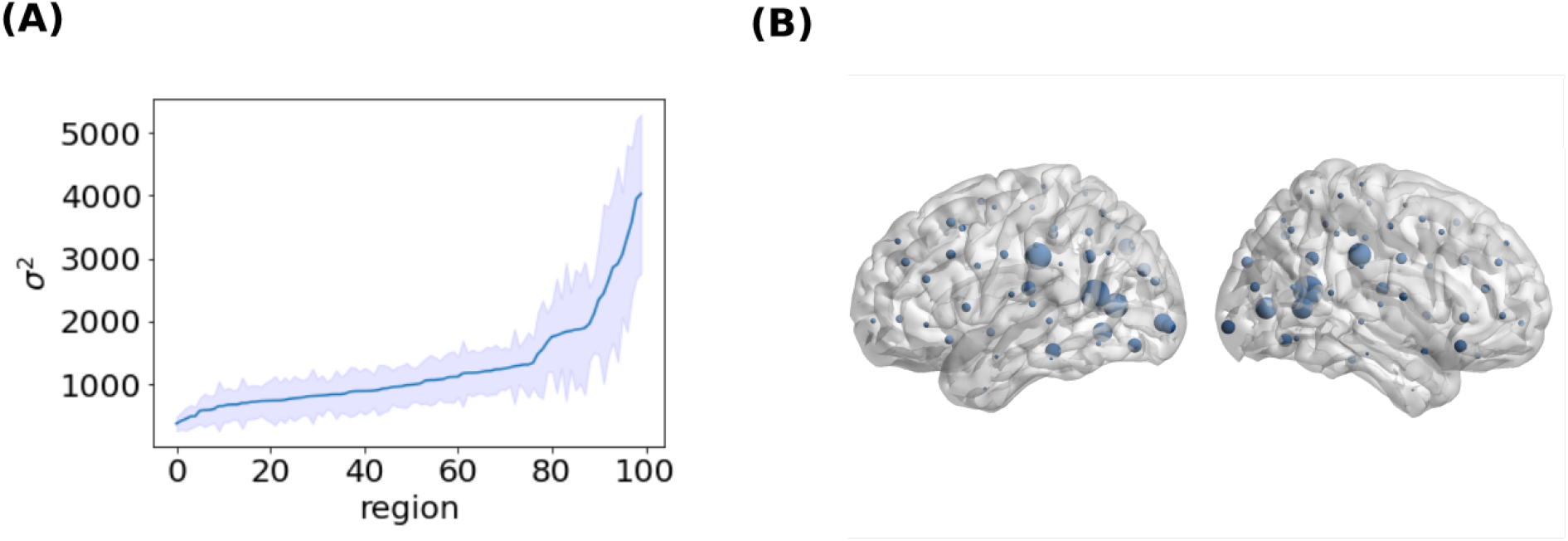
Noise affecting different nodes. **(A)** For each region, we computed the average (over subjects) noise *σ*^2^ and sorted regions by this value. The blue line represents the average (over subjects) noise *σ*^2^, the shaded area represents standard deviation (over subjects). **(B)** In this rendering, the node size is proportional to the average (over subjects) noise *σ*^2^.

**Figure S5:**
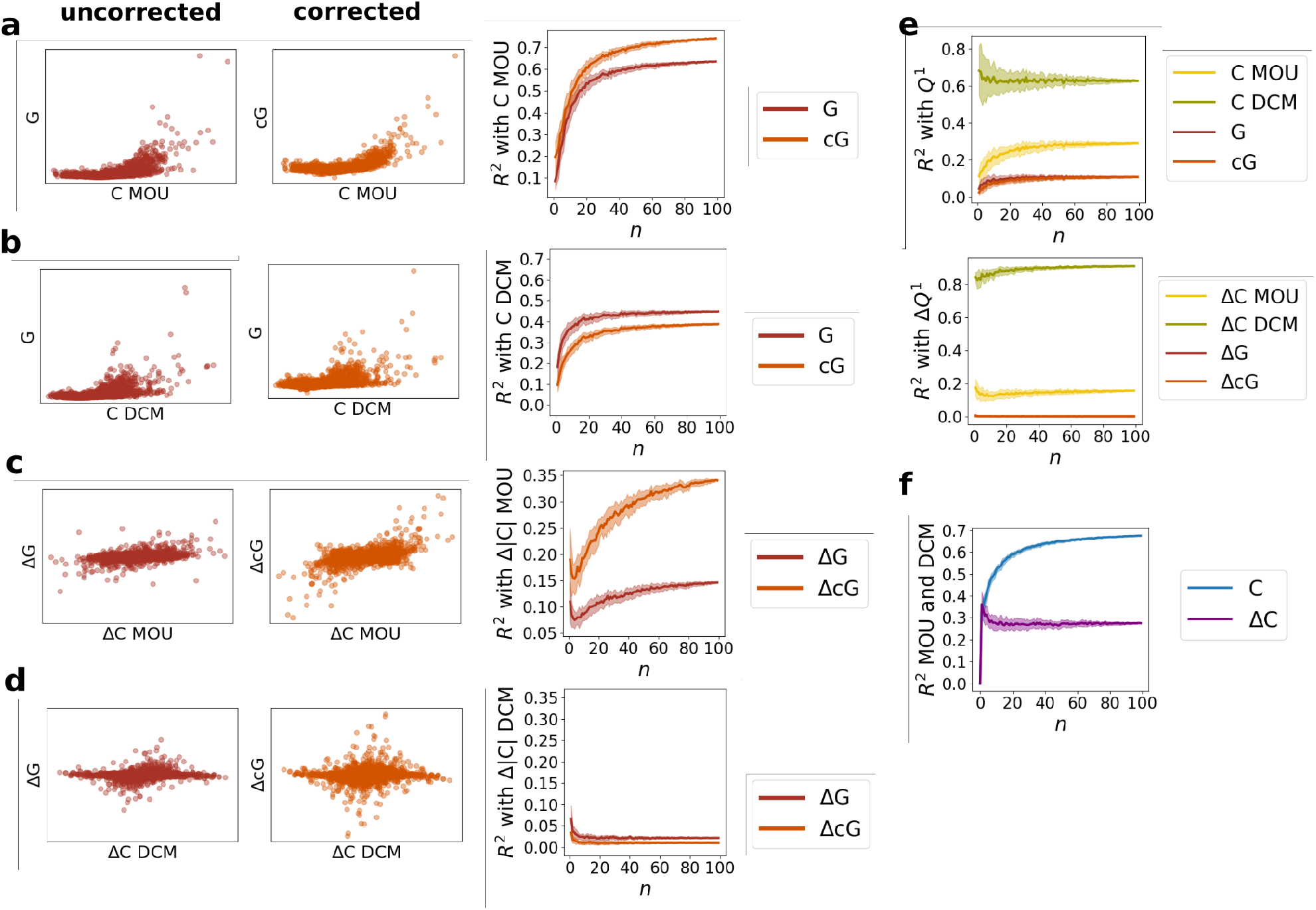
Replication of fig. 4 with independent recording sessions **(A)** effective connectivity *Ĉ* (group average over 100 subjects) vs corrected Granger causality 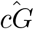 (group average). **(B)** squared Pearson correlation *R*^2^ between group *Ĉ* and group 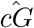 for groups of increasing size *n*. **(C)** effective connectivity *Ĉ* (group average over 100 subjects) vs corrected instantaneous Granger causality 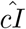 (group average). **(D)** squared Pearson correlation *R*^2^ between group *Ĉ* and group 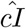 for groups of increasing size *n*. **(E)** asymmetry in effective connectivity Δ*Ĉ* (group average over 100 subjects) vs asymmetry in corrected Granger causality 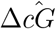 (group average). **(F)** squared Pearson correlation *R*^2^ between group Δ*Ĉ* and group 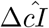 for groups of increasing size *n*.

